# Kinesin-2 from *C. reinhardtii* is an atypically fast and auto-inhibited motor that is activated by heterotrimerization for intraflagellar transport

**DOI:** 10.1101/855940

**Authors:** Punam Sonar, Wiphu Youyen, Augustine Cleetus, Pattipong Wisanpitayakorn, Iman S. Mousavi, Willi L. Stepp, William O. Hancock, Erkan Tüzel, Zeynep Ökten

**Affiliations:** Physik Department E22, Technische Universität München, Garching, Germany; Department of Physics, Worcester Polytechnic Institute, 100 Institute Road, Worcester, Massachusetts; Bioengineering Department, College of Engineering, Temple University, Philadelphia, Pennsylvania; Department of Biomedical Engineering, Pennsylvania State University, University Park, PA 16802, USA

**Keywords:** Intraflagellar Transport, cilia, flagella, *C. reinhardtii*, *C. elegans*, motor cooperation

## Abstract

The construction and function of virtually all cilia require the universally conserved process of Intraflagellar Transport (IFT) [1, 2]. During the atypically fast IFT in the green alga *C. reinhardtii*, up to ten kinesin-2 motors ‘line up’ in a tight assembly on the trains [3], provoking the question of how these motors coordinate their action to ensure smooth and fast transport along the flagellum without standing in each other’s way. Here, we show that the heterodimeric FLA8/10 kinesin-2 alone is responsible for the atypically fast IFT in *C. reinhardtii*. Notably, in single-molecule studies, FLA8/10 moved at speeds matching those of *in vivo* IFT [4], but additionally displayed a slow velocity distribution, indicative of auto-inhibition. Addition of the KAP subunit to generate the heterotrimeric FLA8/10/KAP relieved this inhibition, thus providing a mechanistic rationale for heterotrimerization with the KAP subunit in fully activating FLA8/10 for IFT *in vivo*. Finally, we link fast FLA8/10 and slow KLP11/20 kinesin-2 from *C. reinhardtii* and *C. elegans* through a DNA tether to understand the molecular underpinnings of motor coordination during IFT *in vivo*. For motor pairs from both species, the co-transport velocities very nearly matched the single-molecule velocities, and the complexes both spent roughly 80% of the time with only one of the two motors attached to the microtubule. Thus, irrespective of phylogeny and kinetic properties, kinesin-2 motors prefer to work alone without sacrificing efficiency. Our findings thus offer a simple mechanism for how efficient IFT is achieved across diverse organisms despite being carried out by motors with different properties.

---

Cilia or flagella (used interchangeably) are microtubule-based organelles that project from the surface of most eukaryotic cells. As evidenced from their remarkably diverse deployment across eukaryotes, they are among the most evolutionarily adaptive of all organelles. This diversity is manifested as the large number of seemingly unrelated human disorders linked to impaired ciliary biogenesis, ranging from infertility to vision degeneration [5–12]. Such functional diversity starkly contrasts with the highly conserved nature of IFT, the process that builds and maintains virtually all cilia from unicellular organisms up to humans. IFT was first observed in the unicellular green alga *C. reinhardtii* as movement of large, non-membrane-bound ‘trains’ between the ciliary tip and base (Figure 1) [13]. Given that these trains moved on axonemes – an elaborate microtubule-based scaffold – it soon became clear that IFT is powered by cilia-specific kinesin-2 and dynein-2 motors [1, 14–21]. The kinesin-2 motors that carry out IFT appear to have co-evolved with the axonemal structure, suggesting that their mechanochemical properties are adapted to the highly specialized ciliary environment [22, 23]. Unlike most kinesins that form homodimers, the canonical ciliary kinesin-2 FLA8/FLA10/KAP in *C. reinhardtii* is a heterotrimeric complex consisting of two distinct motor subunits and an accessory, non-motor subunit [4, 14, 24, 25]. Indeed, heterotrimerization of the kinesin-2 motor with the KAP subunit proved to be a universally required property of IFT from the unicellular *C. reinhardtii* up to mammals [23, 25–28]. KAP is suggested to serve as a cargo adaptor for the kinesin-2 motor, however, the physiological relevance of heterotrimerization remains unclear [23].

**Figure 1:**
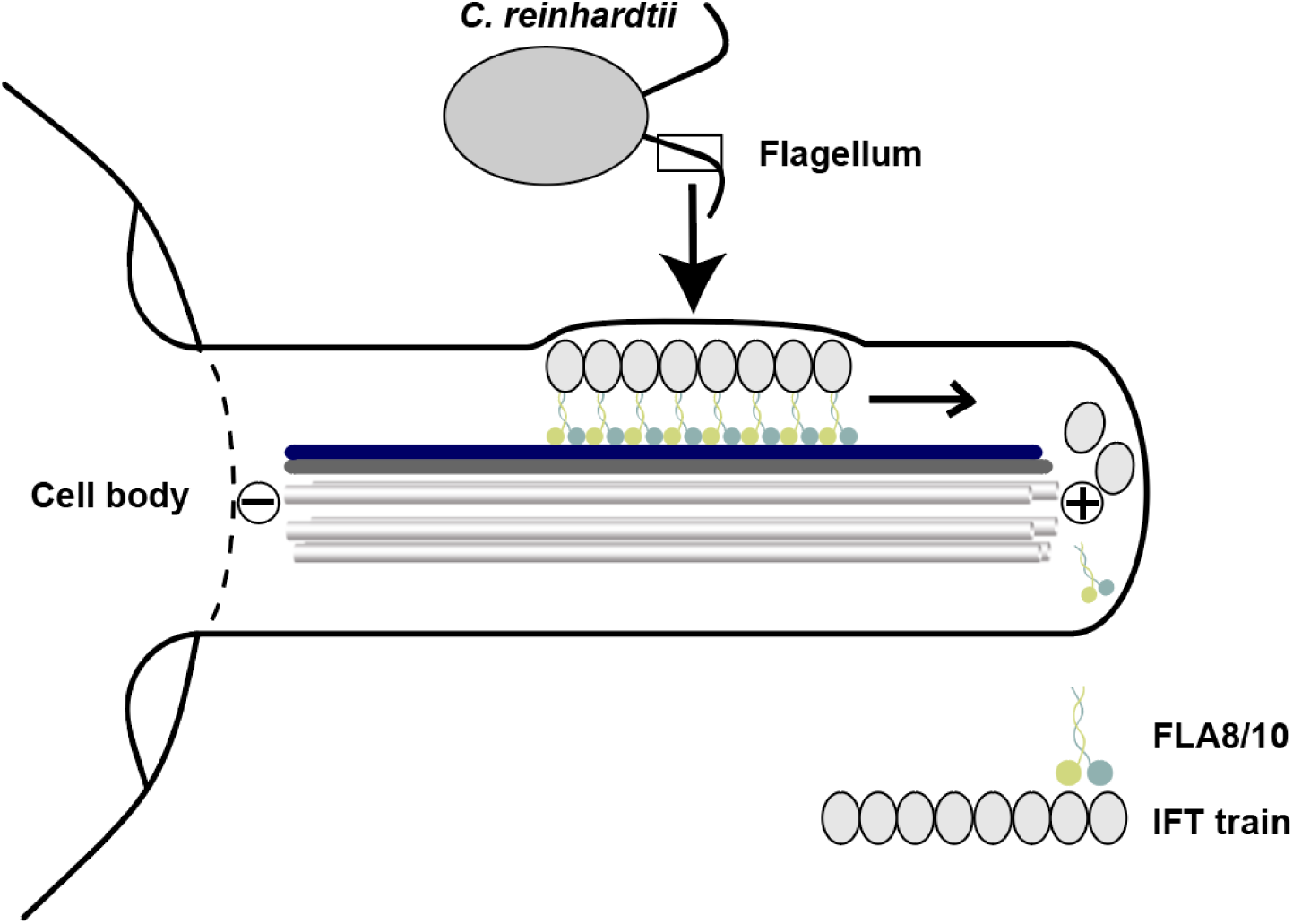
Schematics of kinesin-2-driven IFT in *C. reinhardtii*. Linear arrays of IFT particles or ‘trains’ are transported towards the plus-end of the axoneme. This kinesin-2-driven anterograde transport takes place on the B-tubule of the microtubule doublet (highlighted in blue), leaving the A-tubule (highlighted in dark gray) for the dynein-2 driven IFT towards the minus-end of the axonemes (not shown).

Curiously, although kinesin-2-driven IFT trains in *C. reinhardtii* move with a velocity of ∼2000 nm/s [4], all subsequently characterized heteromeric kinesin-2 motors from diverse model organisms move roughly four-fold slower, both *in vivo* and *in vitro* [27, 29–34]. What is not known is whether FLA8/10 motors display such atypically high velocities in isolation, or whether the fast IFT speeds result from cooperative effects of the assembled motor groups in the *C. reinhardtii* model. Consistent with this notion, functional studies with the *C. reinhardtii* flagellum estimated that up to ten kinesin-2 and dynein-2 motors are reciprocally engaged to move the IFT trains either towards the tip or the base of the cilium, respectively [3]. Given an average train size of ∼200 nm [35], transport toward the ciliary tip thus involves up to 20 active kinesin-2 head domains that ‘line up’ along the train with close spacing between dimers. Moreover, these kinesin-2 motors were shown to exclusively use the B-tubule of the microtubule-doublet of the axoneme, which prevents head-on collisions with retrograde trains moving along the A-tubulin [35]. Such specialized transport geometry raises the question of how the assembled head domains on each train coordinate their actions to achieve efficient IFT *in vivo* (Figure 1).

In contrast to the kinesin/dynein-driven bi-directional transport of cargo in the cytoplasm, which displays frequent reversals, IFT trains move without interruption, out to the ciliary tip via kinesin-2, or back to the base via dynein-2 [2, 36, 37]. This behavior suggests that the oppositely directed kinesin-2 and dynein-2 motors are activated reciprocally during IFT. In support, recent cryo-EM studies with the *C. reinhardtii* flagellum provide compelling evidence that kinesin-2-driven IFT trains transport the minus-end directed dynein-2 as inactive cargo towards the ciliary tip [38]. The underlying molecular mechanisms of how kinesin-2 and dynein-2 motors are specifically activated and inhibited during IFT remain largely unknown. It is conceivable that heterodimeric kinesin-2 also requires an adaptor to be recruited and activated for IFT, as previously demonstrated with the homodimeric OSM-3 kinesin-2 from the *C. elegans* model [39].

Here we turned to *in vitro* reconstitution assays and *in silico* simulations to address following questions. Is the heterodimeric FLA8/10 motor from *C. reinhardtii* alone sufficient to explain the atypically fast IFT velocity of 2000 nm/s *in vivo* [13], or does the fast speed result from the collective motor behaviors? Second, how do populations of FLA8/10 motors coordinate their actions to achieve the smooth and continuous transport observed *in vivo*? Lastly, how are FLA8/10 motors activated and inhibited during IFT?

To gain a mechanistic understanding of *C. reinhardtii* IFT, we recombinantly expressed the FLA8 and FLA10 subunits of the heterotrimeric kinesin-2. The Flag-tagged FLA8 subunit robustly co-precipitated the 6XHis-tagged FLA10, demonstrating the specific heterodimerization between the two different motor subunits (Figure S1A). Next, we turned to functional single-molecule and multiple-motor filament gliding assays to investigate whether a single FLA8/10 motor can move processively on surface-attached microtubules, and if so, whether the single motor speeds differ from the speed of motors working in ensembles.

To this end, we labeled the SNAP-tag on the FLA10 subunit with a fluorescent dye and tracked the movement of single fluorescently labeled motors (Figure 2A). The heterodimeric FLA8/10 motors not only moved processively but also displayed two velocity populations (Figure 2A). The faster velocity population at ∼2000 nm/s demonstrates that this heterodimeric FLA8/10 motor from *C. reinhardtii* alone is sufficient to achieve the atypically fast kinesin-2-dependent IFT velocities observed *in vivo* [13]. The slower velocity population, centered around 1000 nm/s, suggests that, like other full-length kinesins [4, 40–44], this motor is capable of auto-inhibition.

**Figure 2:**
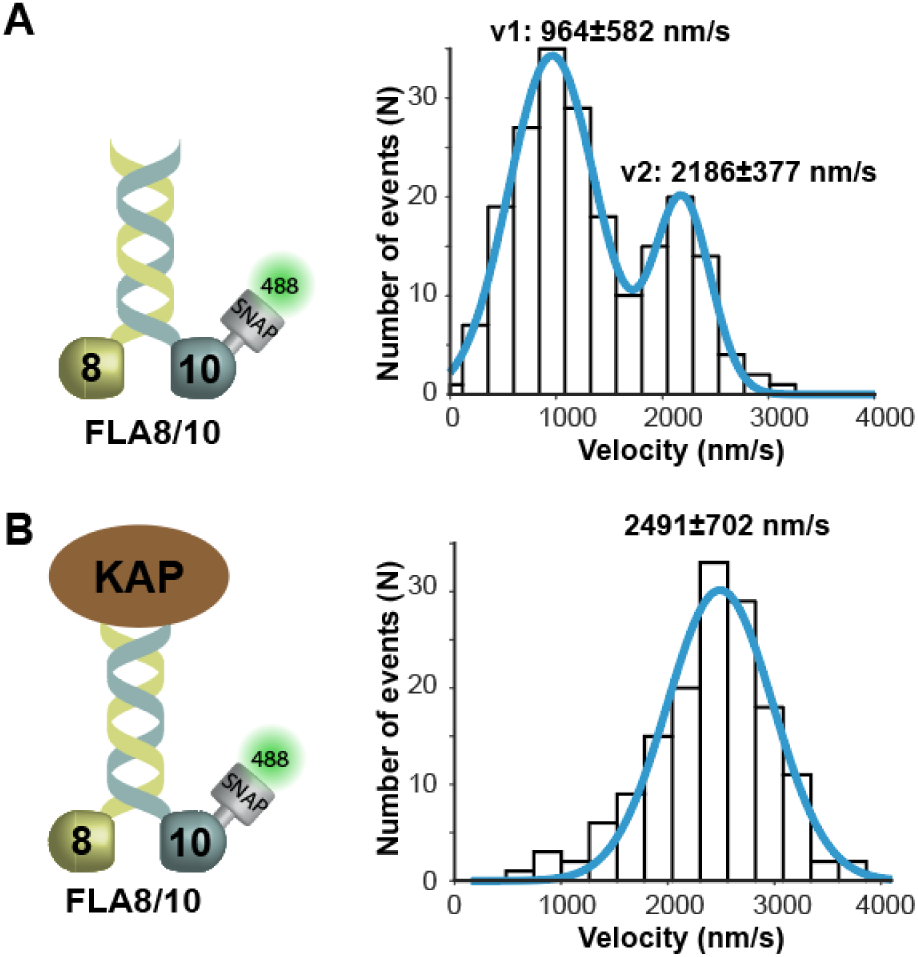
Full-length heterodimeric FLA8/10 is an atypically fast and auto-inhibited kinesin-2 motor. **(A)** Left panel illustrates the wild type FLA8/10 motor that was tracked via the fluorescently labeled N-terminal SNAP-tag on the FLA10 subunit. Single motors displayed two distinct velocity distributions (v1= 964±582 nm/s and v2= 2186±377 nm/s (N=202)) with the fast v2 corresponding to the *in vivo* anterograde transport velocity in *C. reinhardtii* [13] and the slow v1 to auto-inhibition of the wild type FLA8/10 motor, respectively. **(B)** Left panel depicts the heterotrimeric FLA8/10/KAP complex that was tracked via fluorescently labeled SNAP-tag on the N-terminus of the FLA10 subunit. Addition of the KAP subunit abolished the slow velocity distribution (v1) that is observed with the wild type FLA8/10 (A), and the heterotrimeric FLA8/10/KAP complex displayed a fast velocity of 2491±702 nm/s (N=151). Data is fitted to a Gaussian distribution (±width of distribution, from two independent experiments).

In other kinesins, folding of the distal C-terminal ‘tail’ domain onto the N-terminal catalytic heads suppresses the ATPase activity of the dimeric motor, and attaching motors to a surface via their C-terminal tail domains in multiple-motor filament gliding assays relieves this inhibition [29, 30]. Consistent with this, we observed consistently fast velocities in multiple-motor assays (Figure S2A, left, Suppl. movie 1). Thus, the atypically fast IFT velocities in *C. reinhardtii* do not result from collective motor behaviors, but instead are inherent and unique to the FLA8/10 kinesin-2 motor (Figure 2A vs Figure S2A, left). Additionally, a subset of our recombinant full-length FLA8/10 heterodimers are auto-inhibited, likely through autoinhibitory folding of the tail domains onto the head domains.

Intrigued by the sub-population of inhibited motors (Figure 2A), we next asked whether heterotrimerization with KAP may directly alter the activation state of FLA8/10. Previous *in vivo* investigations with *C. reinhardtii* showed that the KAP subunit is required both for achieving efficient processive movement of anterograde IFT trains, and for proper targeting of the kinesin-2 motor to the ciliary base [25]. Consistent with this, assembling the motor subunits into the heterotrimeric FLA8/10/KAP complex (Figure S1B) completely abolished the auto-inhibition, and resulted in all motor complexes moving at full speed (Figures 2A vs 2B). The ability of the KAP subunit to activate the FLA8/10 motor provides a first mechanistic rationale for the assembly of the physiologically-relevant heterotrimeric motor complex *in vivo* and underscores the importance of KAP in the efficient activation of the FLA8/10 motor. The role of the KAP subunit in recruiting [25] and activating *C. reinhardtii* FLA8/10 (Figure 2B) is conceptually highly similar to the adaptor-dependent recruitment and activation of the homodimeric kinesin-2, OSM-3, that carries out IFT in *C. elegans* [39, 45].

Because catalytic head domains are particularly well-conserved across the kinesin-2 family, a mechanistic explanation for the pronounced kinesin-2 speed differences between the single-celled *C. reinhardtii* (∼2000 nm/s) and multicellular organisms (∼500 nm/s) is not immediately obvious [29–31, 33, 34, 46]. Intrigued by these findings, we next asked whether we can delineate the domain responsible for the observed kinetic differences between the respective model organisms. To this end, we replaced the KLP11 and KLP20 head domains in the KLP11/20 heterodimer [29] with the corresponding FLA10 and FLA8 head domains respectively (Figure S2B). Strikingly, this chimeric motor was not only active but also displayed a velocity of >1700 nm/s. Thus, the catalytic heads and not sequences in the coiled-coil or tail domains are the main determinants of the atypically fast kinesin-2-driven IFT velocity in *C. reinhardtii* (Figure S2B).

We next turned to the more complex question of how the multiple kinesin-2 motors that line up in close proximity along each train (Figure 1), cooperate during IFT. To mimic this multi-motor transport *in vitro*, we used double-stranded DNA (dsDNA) to couple two kinesin-2 motors, i.e. four head domains, in close proximity (Figure S3). Given that the FLA8/10 motor is partially auto-inhibited (Figure 2A), we turned to protein engineering to create a fast and fully active kinesin-2 that matches the active anterograde transport motors *in vivo*. To this end, we replaced the FLA10 head with the FLA8 head domain, resulting in FLA8/8 that was heterodimerized by the FLA8 and FLA10 stalks. This chimeric motor no longer displayed the auto-inhibited population seen with the FLA8/10 (Figure 2A), and moved with fast velocities of >2000 nm/s in both single-molecule assays and multiple-motor filament gliding assays (Figure S2A, middle and Figure S2C). In contrast, replacing the FLA8 with the FLA10 head resulted in a chimeric FLA10/10 motor that was considerably slower than the FLA8/8 chimera (Figure S2A, middle vs right), though still substantially faster than IFT kinesin-2 from *C. elegans* and other multicellular organisms [29, 31–34, 46]. Thus, the fast speed of *C. reinhardtii* FLA8/10 cannot be explained by differing activities of the two heads somehow combining to achieve fast transport; instead, the fast speeds are a property of each head individually. *C. elegans* KLP11/20 also combines a faster KLP20 with a slower KLP11 head [29], indicating that, despite being distantly related, the unicellular *C. reinhardtii* and the multicellular *C. elegans* share conceptual similarities. However, possessing different kinetic signatures between the two heterodimerized heads is insufficient to explain the substantial velocity differences between the two model organisms.

To examine whether the fast *C. reinhardtii* and the slow *C. elegans* kinesin-2 IFT motors differ in their abilities to cooperate during multi-motor transport of IFT trains, we compared the co-transport behaviors of the constitutively activated *C. reinhardtii* FLA8/8 (Figure S2C) with that of *C. elegans* KLP11/20 [29]. The heterodimeric nature of kinesin-2 allowed for a well-defined and highly specific coupling and fluorophore labeling of the two motors. Specifically, we introduced a N-terminal SNAP-(for fluorophore labeling) and C-terminal Halo-tag (for dsDNA coupling) into the FLA8 subunit, respectively. The co-expression of this functionalized FLA8 subunit with the chimeric FLA8^head^FLA10^stalk^ motor resulted in a kinesin-2 motor with the N-terminal SNAP- and C-terminal Halo-Tag on *one* subunit only (Figure S3A). Next, we functionalized dsDNA, containing thiol groups at both ends, to the Halo Iodoacetamide O4 ligand, which covalently links to Halo-Tags (Figure S3B and S3C, see Supplementary Information for details). With this strategy, coupled kinesin pairs were generated by covalently linking motors with C-terminal Halo-Tags to each Iodoacetamide-functionalized end of the dsDNA [47, 48]. To ensure that only coupled motor pairs are analyzed, motors were labeled on their N-terminal SNAP-tag with different fluorophores as illustrated in Figure S3A. Simply mixing the two differentially labeled FLA8/8 motors with the bi-functionalized dsDNA resulted in motor-DNA hybrids (FLA8/8-FLA8/8 hereafter) that could be distinguished in *in vitro* reconstitution assays by the movement of co-localized green and red fluorophores (Figure S3D, Suppl. movie 2). Using the same strategy, we also coupled two constitutively activated KLP11/20 motors from *C. elegans* for direct comparison [29] (KLP11/20-KLP11/20 hereafter) (Figure S3E, Suppl. movie 3). Collectively, the heterodimeric nature of kinesin-2 allowed a highly stable and specific DNA-motor coupling and fluorescent labeling with the respective functionalization being unambiguously assigned to the subunits (Figures S4 and S5).

To gain mechanistic insight into the forces generated and the degree of inter-motor coordination during co-transport by FLA8/8 and KLP11/20 motor pairs, we developed coarse-grained simulations of motor stepping, described fully in Materials and Methods. Briefly, motors are modeled as stochastic steppers and the motor-DNA complex is modeled mechanically as a Freely Jointed Chain (FJC), with segments corresponding to distinct sub-domains of the complex (Figure 3A). The two dimeric motors bind and unbind from the microtubule at specified rates and when one motor detaches from the microtubule, it is subjected to Brownian (Figure S6A) and entropic restoring forces from the FJC linkage (Figure S6B). Motors step along the microtubule with a linear force-velocity profiles and a 6 pN stall force (Figure S7A). Unloaded motor detachment rates were calculated from experimental single-molecule run lengths and velocities summarized in Tables S1 and S2. In this model, FLA8/8 and KLP11/20 are assumed to have identical force-dependent detachment rate profiles (*k*_off_(*F*)) at F>2 pN and F<-1 pN as their kinesin-2 peer, KIF3A/B, which has been thoroughly characterized by optical trapping experiments [46]. As no experimental data are available for force-dependent detachment rates at very low forces, exponential interpolations were used to connect detachment rates at -1 pN and +2 pN to the measured zero-load motor off-rate (Figure S7B; see Materials and Methods for further details). Using this approach, simulated single-molecule velocities of FLA8/8 (1954 ± 117 nm/s) and KLP11/20 (587 ± 82 nm/s; N=1000 runs for each), calculated by linear fits to position vs time traces (Figure 3B and Figure S8A), were in good agreement with experimental values (Table S1). The corresponding run length distributions for FLA8/8 (Figure 3C; mean 2.12 ± 0.13 μm) and KLP11/20 (Figure S8B; mean 2.28 ± 0.15 μm) also agreed well with experiments (Table S1).

**Figure 3:**
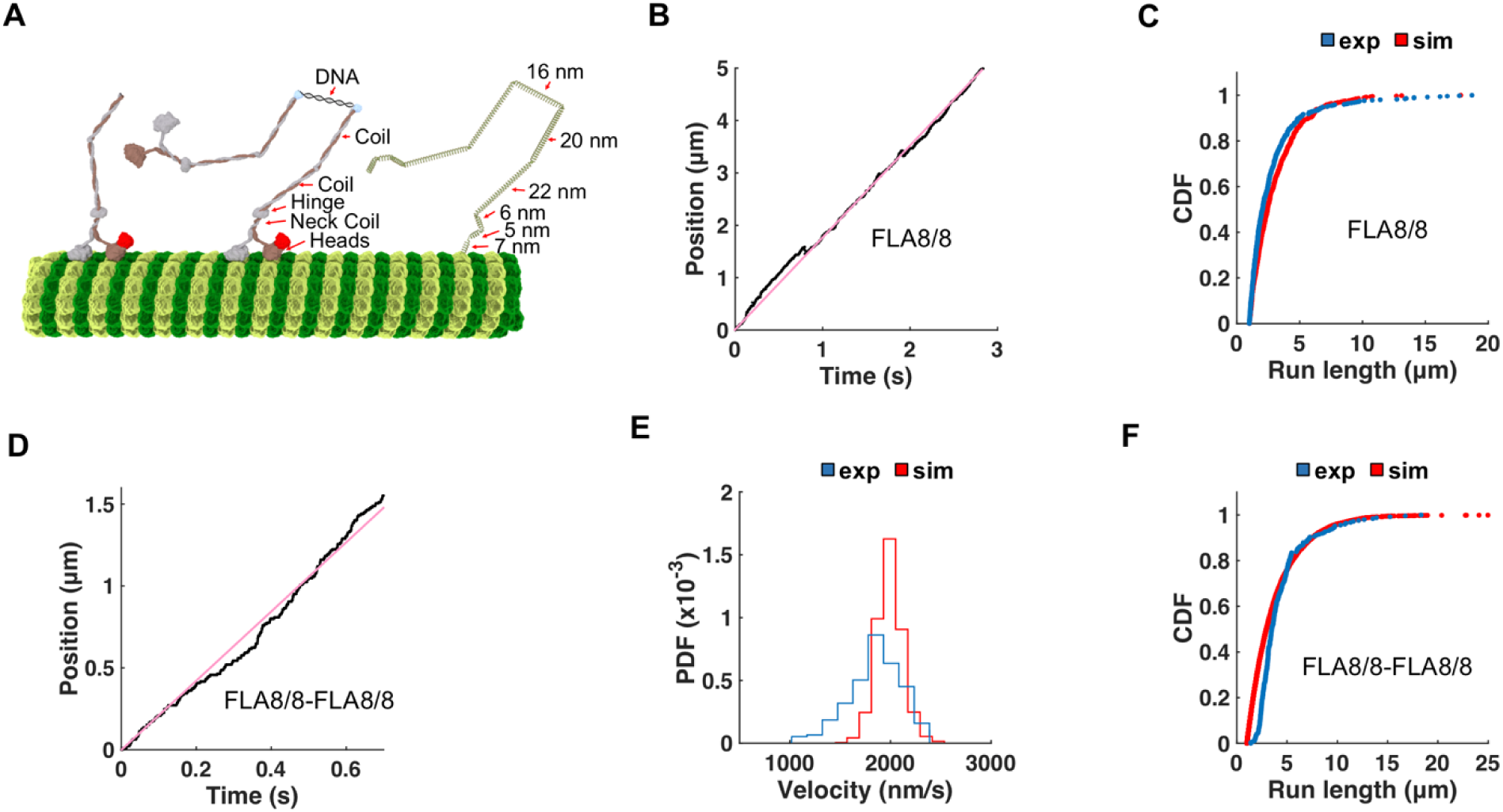
Model recapitulates kinetic parameters of FLA8/8 and FLA8/8-FLA8/8 co-transport. **(A)** Illustrations of a single kinesin-2 (left), a motor-DNA hybrid construct (middle), and the FJC simulation model (right). The springs shown on the right correspond to the two kinesin-2 motors (head, neck coil, hinge, coil1, coil2), and the dsDNA. **(B)** Positions (black) of a single FLA8/8 motor from the simulations as a function of time (tracked at the head). The simulated velocities were calculated from the slope of each trace (pink line). **(C)** Cumulative distributions of experimental (blue) and simulated (red) run lengths of a single FLA8/8 motor (Table S1). **(D)** Positions (black) of the FLA8/8-FLA8/8 co-transport from the simulations as a function of time (tracked in the middle of the dsDNA). **(E)** Experimental (blue) and simulated (red) velocity distributions of FLA8/8-FLA8/8 co-transport. The means and standard deviations were determined by fitting to a Gaussian function; 1806 ± 339 nm/s (experiment), 1927 ± 124 nm (simulation). The simulated velocity was calculated from the slope of each trace (pink line). **(F)** Cumulative distributions of experimental (blue) and simulated (red) run lengths of FLA8/8-FLA8/8. The mean run lengths determined by fitting the cumulative distribution function (CDF) are 3.00 ± 0.06 μm for the experiment and 2.80 ± 0.11 μm for the simulation, respectively.

Having validated our single-molecule simulations, we next turned to modeling the co-transport behaviors of FLA8/8-FLA8/8 (Figure 4A) and KLP11/20-KLP11/20 (Figure 4B). Simulations were initialized by binding of one motor to the microtubule and allowing the second motor to attach with a reattachment rate (*k_on_*) that was the only open parameter in the simulations. Simulations terminated when both motors detached from the microtubule. The motor-DNA hybrid was tracked at the motor domains when only one motor was bound, and at the mid-point between the two motors when both motors were bound. Sample traces of simulated FLA8/8-FLA8/8 and KLP11/20-KLP11/20 co-transport are shown in Figure 3D and Figure S8C respectivaly. Reattachment rates *k_on_* were varied to find the best match of simulations to the experimental velocities and run lengths of the corresponding motor pairs. The reattachment rates that best recapitulated the experiments were 4 s^-1^ for FLA8/8 and 2.5 s^-1^ for KLP11/20 (Figures 3E and 3F, Figures 4C–G, Figures S8D and S8E, Figure S9, Suppl movies 4 and 5). The velocity and run length values are summarized in Table S1, and simulation parameters are given in Table S2.

**Figure 4:**
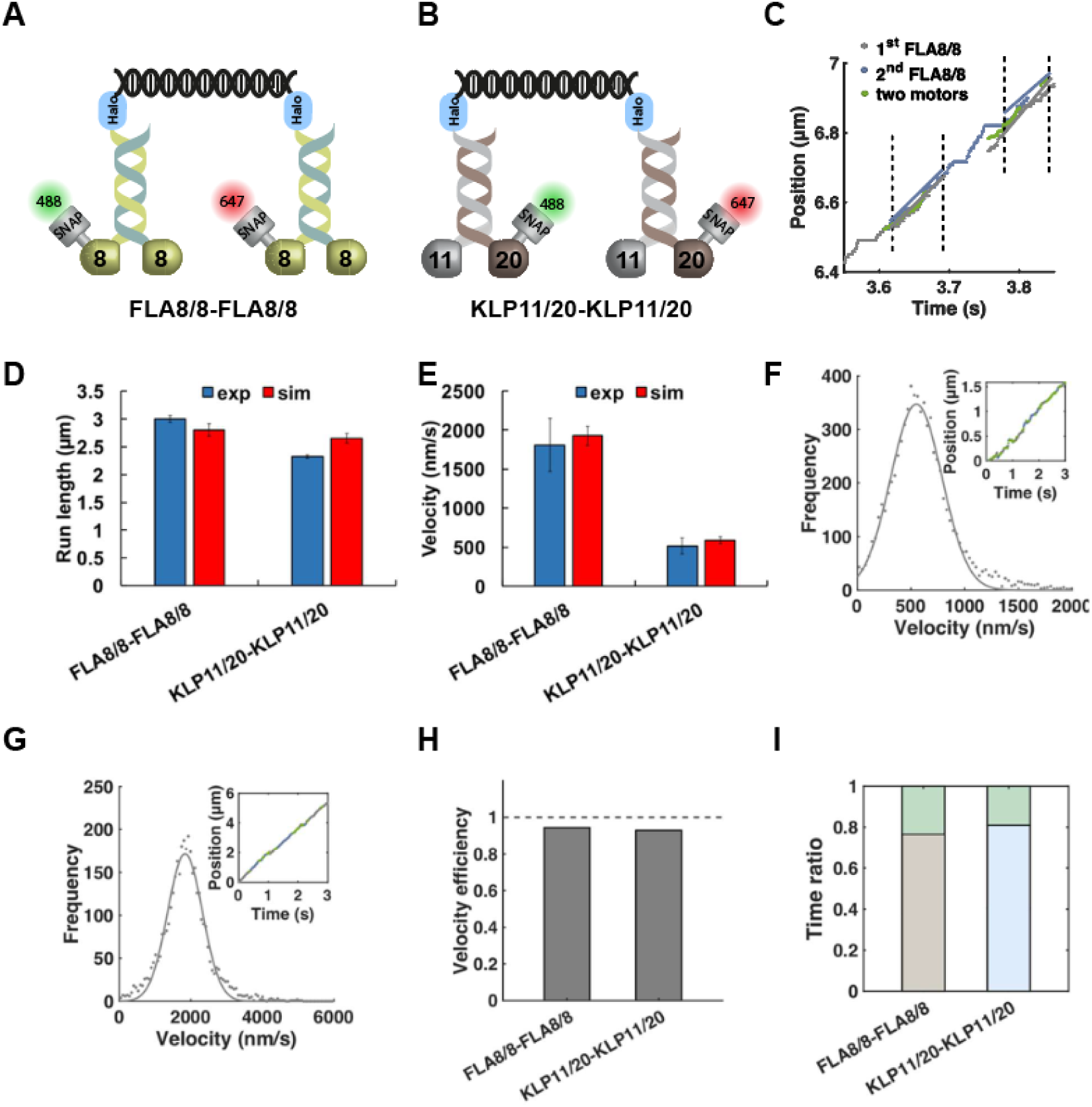
Simulations recapitulate experimental results of co-transport by FLA8/8 and KLP11/20 pairs. (**A) and (B)** depict coupling of two fast and two slow motors, respectively. **(C)** Simulated trace of FLA8/8-FLA8/8 co-transport. Trace shows position of the first motor (grey) or the second motor (blue) when only one motor is bound to the microtubule, or the position of the middle of the complex (green) when both motors are simultaneously bound to the microtubule. The regions bounded by two vertical dashed lines represent periods of co-transport. **(D)** Run length, and **(E)** velocity data comparing experiments (blue) with simulations (red). Error bars indicate mean ± SD. **(F) and (G)** Velocity histogram of KLP11/20 **(F)** and FLA8/8 **(G)** motor-DNA hybrids during their co-transport. Co-transport velocities (grey circles) in simulations were measured by linear fits to the traces during periods when both motors were bound to the microtubule; see (C) for FLA8/8-FLA8/8 co-transport and Figure S9 for KLP11/20-KLP11/20 co-transport. **(H)** Motor velocity efficiencies (co-transport velocity divided by the unloaded velocity of single motors) during co-transport for FLA8/8-FLA8/8 (0.94 efficiency) and KLP11/20-KLP11/20 (0.93 efficiency). **(I)** Fraction of time simulated motor-DNA hybrids spent in one- or two-motor bound states. Of total transport time, FLA8/8-FLA8/8 spent 0.76 with one motor bound (gray), and KLP11/20-KLP11/20 spent 0.81 of time with one motor bound (blue).

Having identified a parameter set that recapitulates the experimental behavior of the motor pairs, we then used the simulations to investigate motor behavior and inter-motor coupling during co-transport. To determine whether different motors are able to better coordinate in groups than others, we calculated the co-transport velocity efficiencies by dividing the average velocity of the motors during co-transport by their respective unloaded velocities. Based on the simulations, the velocity efficiencies of FLA8/8 and KLP11/20 were similar and were both only slightly lower than 1 (Figure. 4H); thus, by this metric their co-transport abilities were similar despite their four-fold different speeds. Next, to investigate the attachment/detachment dynamics during co-transport, we calculated the fraction of time each complex spent with only one motor bound versus both motors bound (Figure 4I). For both FLA8/8-FLA8/8 and KLP11/20-KLP11/20, the complexes spent roughly 80% of the time with only one motor bound and transporting the complex. Thus, in the majority of traces, the second motor could not land before the first motor detached from the microtubule, or one of the motors detached due to the high detachment rates under load during co-transport. This result is consistent with the finding that the run lengths of the FLA8/8-FLA8/8 and KLP11/20-KLP11/20 complexes were only slightly higher than their corresponding single-molecule run lengths (Table S1). To summarize, when working in pairs, both FLA8/8 and KLP11/20 generally work as single-molecules rather than cooperating with their partner in the complex.

Collectively, our *in vitro* and *in silico* dissections provide key mechanistic insights into IFT in the flagella of *C. reinhardtii* where IFT by kinesin-2 was first identified [13]. We demonstrate that heterotrimerization with the KAP subunit, as occurs *in vivo*, relieves auto-inhibition in the heterodimeric FLA8/10 kinesin-2 and fully activates the motor for efficient and atypically fast processive transport *in vitro*. The heterotrimeric FLA8/10/KAP transport complex is thus necessary and sufficient for the previously observed anterograde IFT in *C. reinhardtii*. We pinpoint this so far unique kinetic property to the catalytic heads of the heterodimeric FLA8/10. However, irrespective of species-specific kinetics, the kinesin-2 motors from the unicellular *C. reinhardtii* and multicellular *C. elegans* both display similar uncooperative behavior during co-transport suggesting that these motors share common principles to accomplish efficient IFT *in vivo*.

## Acknowledgments

This work was supported by the European Research Council Grant (335623) to Z.Ö. and National Institute of Health Grants No. R01GM100076 and No. R01GM121679 to W.O.H and E.T. We acknowledge the Thai government for supporting W.Y. and P.W. through the Development and Promotion of Science and Technology (DPST) Scholarship. The authors would also like to thank all members of the Ökten and Tüzel groups for helpful discussions and their insightful suggestions.

## Author contributions

P.S. and Z.Ö. designed the experiments. P.S. and A.C. performed experiments and analyzed the data. W.L.S. wrote all customized MATLAB routines. W.Y. developed the model, performed the simulations, analyzed data, prepared the figures; P.W. analyzed data, prepared figures; I.S.M. developed the Monte Carlo simulations for calculating the landing distributions; E.T. supervised all the modeling work. Z.Ö., P. S., W. O. H, W.Y., P.W. and E.T. contributed to the manuscript writing.

## Competing financial interests

The authors declare no competing financial interests.

## Supporting information

### SI Materials and Methods

All reagents were the highest purity available and were obtained from Sigma-Aldrich unless stated otherwise.

#### Protein expression, purification and fluorescent labeling

Proteins were expressed using the Baculovirus Expression System in insect cells (Spodoptera frugiperda, Sf9) according to the manufacturer’s instructions (Life Science Technologies). Proteins were Flag-tagged (DYKDDDDK) at their C- or N-terminal ends to facilitate purification. Cells (protocol for 50 ml culture) were harvested by centrifugation @1500 rpm for 15 minutes. Purification was performed at 4 °C. Cell pellets were lysed in lysis buffer (10% Glycerol, 50 mM Pipes, pH 6.9, 300 mM potassium acetate (KAc), 1 mM MgCl_2_, 1 mM DTT, 0.1 mM ATP, 0.5% Triton X-100, complete protease inhibitor tablet (Roche)). Cell debris were removed by centrifugation at 30000 rpm for 10 minutes at 4°C. Supernatant was incubated with 100 µl of Anti-Flag M2 affinity gel at 4 °C for 90 minutes. Beads were centrifuged at 800 rpm for 15 minutes at 4 °C and washed three times with 1 ml of wash buffer 1 (10% Glycerol, 80 mM Pipes, pH 6.9, 500 mM KAc, 1 mM MgCl_2_, 1 mM DTT, 0.1 mM ATP, 0.1% Tween-20, complete EDTA-free protease inhibitor cocktail (Roche)) and three times with 1 ml of wash buffer 2 (10% Glycerol, 80 mM Pipes, pH 6.9, 200 mM KAc, 1 mM MgCl_2_, 1 mM EGTA, 1 mM DTT, 0.1 mM ATP, 0.1 % Tween-20, complete EDTA-free protease inhibitor cocktail (Roche)). For fluorescent labeling, beads were incubated for 45 minutes on a rotator at room temperature with wash buffer 2, containing 10 µM SNAP-Surface Alexa Fluor^488^ or SNAP-surface Alexa Fluor^647^ (New England Biolabs). Excess dye was removed by washing the beads three times with 1 ml of wash buffer 2, and labeled protein was eluted in elution buffer (Wash buffer 2 containing 0.5 mg/ml 1X Flag peptide) after 45 minutes of incubation at 4 °C. Purified proteins were analyzed by SDS-PAGE on a 10% polyacrylamide gel. Multi-motor gliding assays were performed as described previously to ensure motor protein quality [29].

#### Proteins used in this study

1. FLA8^C-Flag^/FLA10^C-His^
2. FLA8^C-Flag/ N-SNAP^FLA10^C-His^
3. ^N-SNAP^FLA8^C-Halo-Flag^/FLA8^(10 stalk)C-His^
4. FLA10^(8 stalk) C-Flag^/FLA10^C-His^
5. KLP11^C-Halo-His/N-SNAP-Flag^KLP20 (with activating mutations as described in [29])
6. FLA8^(KLP20 stalk) C-Flag/N-SNAP^FLA^10(KLP11 stalk) C-His^

Splice sites used in this study are indicated below [29].

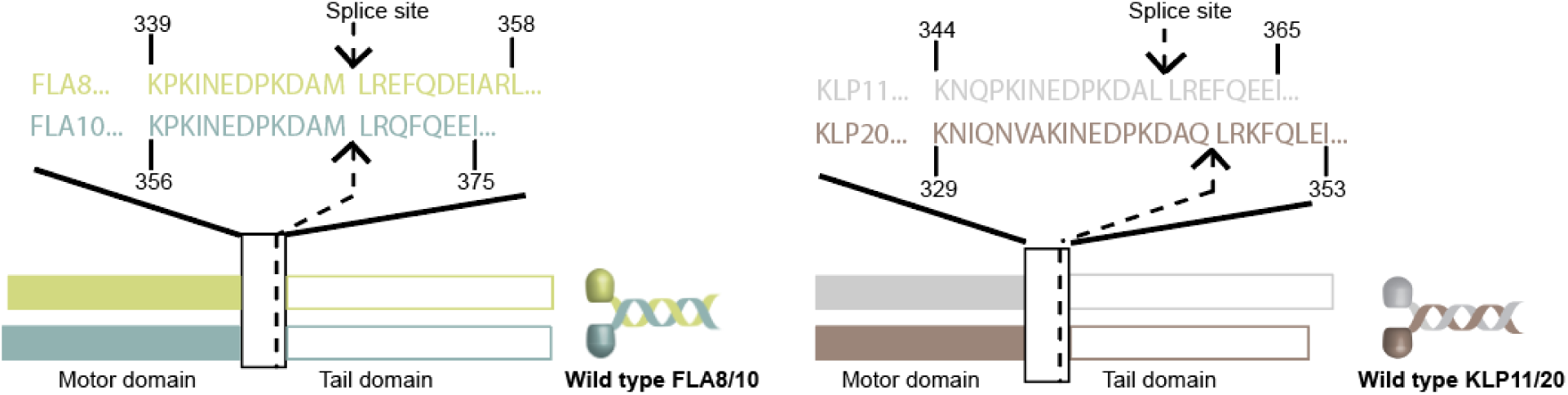

#### Assembly of the Motor-DNA hybrids

##### I. Covalent coupling of one motor to dsDNA

For specific coupling of motor to DNA, double-stranded DNA (dsDNA) was synthesized with Atto-633 dye on one strand and with a thiol group on the complementary strand (Biomers):

5’-ATTO633-CCG AGG ACT GTC CTC CCG AGT GCG GCT ACG ACG TTA CCC GGG TGA GCA-3’

5’-TGC TCA CCC GGG TAA CGT CGT AGC CGC ACT CGG GAG GAC AGT CCT CCG-Thiol C6-3’

###### Purification and activation of fluorescent thiol-dsDNA

The single-stranded DNA oligonucleotides containing the fluorescent dye and the thiol-group were mixed in an equimolar concentration and incubated for 30 minutes at room temperature. The sulfhydryl-groups (-SH) were regenerated by the addition of 1/10 volume of TCEP and the mixture was incubated for 30 minutes at room temperature. The DNA solution was mixed with of 1/10 volume of sodium acetate (NaAc) and subsequently with 2.5× volume of ice-cold ethanol and incubated for 1 hour at -20°C and centrifuged for efficient precipitation. The DNA pellet was centrifuged at 14000 rpm for 30 minutes at 4°C and washed with 70% ethanol. The pellet was resolubilized in ammonium bicarbonate buffer (containing 1mM TCEP) and immediately reacted with five-fold excess of the Halo Iodoacetamide O4 ligand and incubated for 45 minutes on a rotator at room temperature. Unreacted Iodoacetamide ligand was removed via HPLC and the functionalized DNA was stored at -20°C for long term use. The C-terminally Halo-tagged and fluorescently labeled motors (Figure S3) were covalently linked to the Iodoacetamide-functionalized dsDNA [47, 48] by simply mixing in a 1:1 ratio (for coupling specificity and efficiency see Figures S4 and S5).

##### Buffers and solutions

TECP: 100 mM in ddH_2_0

NaAc: 3 M in ddH_2_0, pH 5.2

Ammonium bicarbonate: 200 mM, pH 8.0

Halo Iodoacetamide O4 ligand (Promega): 100 mM in DMSO

Wash buffer: 80 mM Pipes, pH 6.9, 200 mM KAc, 1 mM MgCl_2_, 1 mM EGTA, 1 mM DTT, 0.1% Tween-20, complete EDTA-free protease inhibitor cocktail.

##### II. Covalent coupling of two motors to dsDNA

To specifically couple two motors to dsDNA, same strategy as above has been followed except, in this case dsDNA is unlabeled and one strand of dsDNA is functionalized with two thiol groups that in turn was covalently linked to the Halo Iodoacetamide O4 ligand (Biomers):

5’-CCG AGG ACT GTC CTC CCG AGT GCG GCT ACG ACG TTA CCC GGG TGA GCA-3’

5’-Thiol C6-TGC TCA CCC GGG TAA CGT CGT AGC CGC ACT CGG GAG GAC AGT CCT CCG-Thiol C6-3’

To couple two differently labeled motors (SNAP^488^ and SNAP^647^), the bi-functionalized unlabeled dsDNA was incubated with both motors in equimolar concentrations for 20 minutes at room temperature (for coupling specificity and efficiency see Figures S4 and S5).

#### Buffers for TIRF assays

BRB10: 10 mM Pipes, pH 6.9, 1 mM EGTA, 2 mM MgCl_2_, 5 mM DTT.

BRB10/BSA wash buffer: BRB10 plus 2 mg/ml Bovine Serum Albumin (BSA).

BRB10 motility buffer: BRB10 plus 0.145 mg ml^−1^ glucose oxidase, 0.0485 mg ml^−1^ catalase, and 20% glucose.

#### Single-molecule and colocalization assays

Single-molecule and colocalization assays were performed as previously described [31, 39]. Briefly, biotinylated microtubules were attached to the glass surface of the flow chamber that was pre-coated with biotinylated BSA and Streptavidin (each 1 mg ml^−1^). Fluorescent motors were diluted in BRB10 supplemented with 8 mM ATP, 0.18 mg ml^−1^ glucose oxidase, 0.06 mg ml^−1^ catalase, and 0.4% glucose. Diluted proteins were incubated with surface-attached microtubules for 2 minutes in the flow chamber and washed with BRB10 buffer supplemented with 2mg/ml BSA. Images of single motors walking on filaments were recorded with a cycle time of 76 ms with an objective-type Leica DMI6000 B TIRF microscope (Leica), equipped with an oil-immersion 100× Plan objective lens (numerical aperture 1.47), and a back-illuminated Andor U897 EMCCD camera (Andor). Excitation was achieved by diode lasers at 488 or 638 nm wavelength, and frames were recorded and analyzed with the AF 6000 software (Leica). Velocities and run lengths were analyzed with custom-written programs implemented in MATLAB (MathWorks, Natick, MA). Runs (distance from start position to the position with the largest distance) with minimum 1 µm were considered as processive (*χ*0 = 1 μm). Histograms of velocities were fitted to a Gaussian distribution, and run-lengths were fitted by a single-exponential model to the empirical Cumulative Distribution Function (CDF) of the measured runs. Runs were included only if they moved smoothly for at least 5 frames and had r^2^ value > 95%.

For colocalization assays, the same procedure was followed except two channels (488 nm and 638 nm) were used for simultaneous observation of both motors. The cycle time for both channels was 195 ms. Colocalized movies were analyzed using custom-written routines in MATLAB (MathWorks, Natick, MA). In order to assign colocalized runs, a penalty score was calculated for all combinations of two runs from the respective channels. The penalty score resulted from the mean distances of tracked positions (pixels, factor 1/3) and the difference in the starting time (frames, factor 1). Long runs were cropped to the length of the shorter runs in order to account for bleaching events. Pairs of runs with a penalty score <10 were considered colocalized and their parameters were taken into account in further analysis as the average values of both runs.

#### Microtubule decoration assays

To determine the coupling efficiency of motor-DNA hybrids, decoration assays were performed by preparing the following mixtures and incubating for 20 minutes at room temperature:

**A.** Labeled motor (^N-SNAP488^FLA8/8 or KLP11/ ^N-SNAP488^20) without C-terminal Halo-tag was mixed in equimolar concentration with Atto-633 dsDNA functionalized at one end with thiol plus Halo Iodoacetamide O4 ligand.

**B.** Labeled motor (^N-SNAP488^FLA8^C-Halo^/8 or KLP11^C-Halo^/ ^N-SNAP488^20) was mixed in equimolar concentration with Atto-633 dsDNA functionalized at one end with thiol minus Halo Iodoacetamide O4 ligand.

**C.** Labeled motor (^N-SNAP488^FLA8^C-Halo^/8 or KLP11^C-Halo^/ ^N-SNAP488^20) was mixed in equimolar concentration with Atto-633 dsDNA functionalized at one end with thiol plus Halo Iodoacetamide O4 ligand.

**D.** Differently labeled motors (^N-SNAP488^FLA8^C-Halo^/8, ^N-SNAP647^FLA8^C-Halo^/8 or KLP11^C-Halo^/^N-SNAP488^20, KLP11^C-Halo^/^N-SNAP647^20) were mixed in equimolar concentration with unlabeled dsDNA functionalized at both ends with thiols plus Halo Iodoacetamide O4 ligand. Unlabeled microtubules were attached to the glass surface of the flow chamber coated with a Biotinyted BSA-Streptavidin (1 mg ml^−1^ each) sandwich. Mixture (A/B/C/D) was diluted to the desired concentration in BRB10 motility buffer and flowed into the chamber. Images were captured at different locations of the surface. Colocalization efficiencies were analyzed using a custom-written MATLAB routine (MathWorks, Natick, MA).

##### Computational Model

The model for the motor-DNA hybrid transport is an extension of our earlier work on kinesin-1 and -3 motors [49]. However, instead of explicit Brownian dynamics of the motors attached to a scaffold, it employs a freely jointed chain (FJC) model for the entire kinesin-scaffold-kinesin construct. The whole construct is modeled in three dimensions, and consists of 11 rigid segments of non-uniform-lengths, as illustrated in Figure 3A. The microtubule (MT) is modeled as a 25 nm diameter cylinder anchored on glass surface with the scaffold-motor complex placed on top.

We developed a separate Monte-Carlo simulation to calculate the distribution of the end-to-end distance of the motor-DNA hybrid, *P(R),* and its standard deviation, *σ,* which is used to determine attachment locations in the 3D geometry (Figure 3A). In this model, a point is first placed on the microtubule, and a second point is generated using random values with uniform distributions between [0,π) and [0, 2π) for the two corresponding spherical angels, *θ* and *ϕ*, respectively. We then continued this procedure to grow the scaffold, one segment at a time, with the segment lengths shown in Figure 3A. Any configuration that intersects the cylindrical microtubule at any location is rejected, and the accepted configurations are projected along the backbone of the microtubule. We then calculated a histogram (Figure S6A), and fit it to a Gaussian to calculate the standard deviation, *σ*, of end of end distances, i.e. the landing distance distribution.

During co-transport, both motors are subject to forces of the same magnitude but in opposite directions. The front motor (toward the plus end) is under a hindering load (negative), whereas the rear motor (toward the minus end) is under an assisting load (positive). The force-extension curve is modeled in a piece-wise manner, as shown in Figure S6B, namely

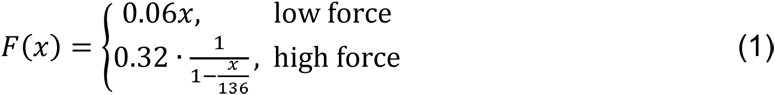

where *x* is the end-to-end distance (nm), *F* is the force acting on the motor (pN), and 136 nm corresponds to the total contour length of the complex.

At each time step, motors stochastically take a step of 8 nm based on a probability of stepping given by

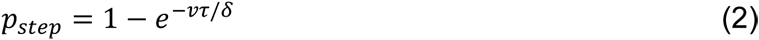

where *v* is the motor velocity, τ is the simulation time step, and *δ* is the step size. The linear force-velocity curve was taken from [50] as follows (see also Figure S7A)

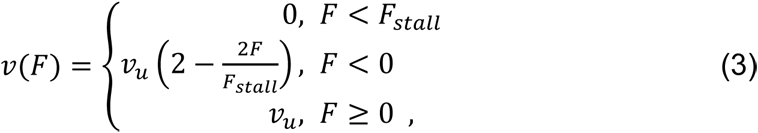

Where *v_u_* is the unloaded velocity, *F* is the force acting on the motor and *F_stall_* is the stall force (6 pN) [29].

Motors stochastically dissociate from the MT based on the force-dependent detachment rate, *k*_off_(*F*), of kinesin-2, previously measured by Milic et al. [46]. Due to the lack of experimental data at low forces (-1 to 2 pN), we utilized exponential interpolations towards the known *k*off,0 value, resulting in the force-dependent detachment rate given by

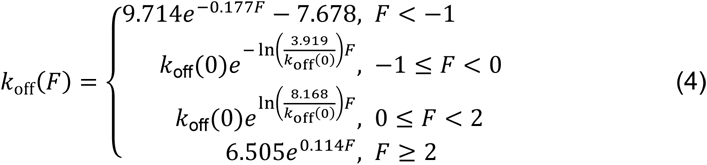

where *k*_off_(0) represents the unloaded detachment rate (= velocity/run length) (plotted in Figure S7B). After a motor detaches from the MT, it diffuses within a range of the contour length (136 nm) and can reattach back to the MT based on the reattachment rate, *k_on_*.

#### Data availability

All raw data sets used for the analysis or (full pictures of decoration images or colocalised movies) and MATLAB codes are available from the corresponding author on request.

## Supplementary Figures

**Figure S1:**
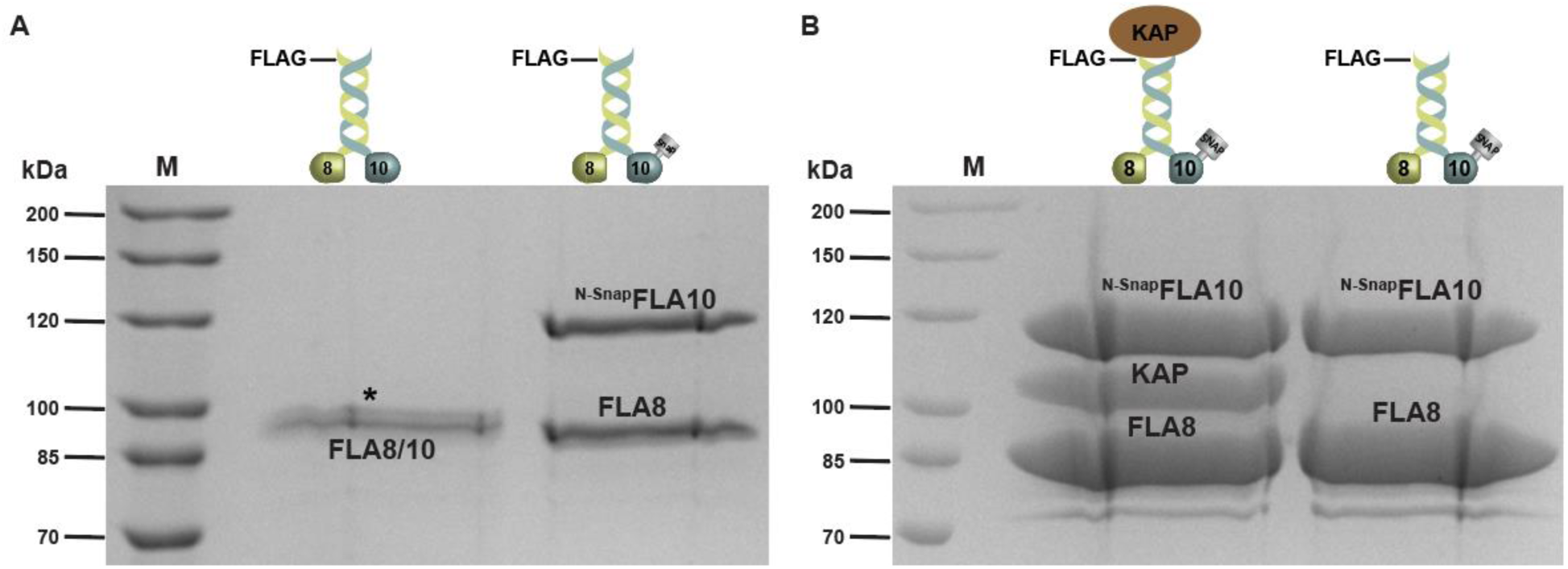
(A) FLA8 and FLA10 subunits form a heterodimeric kinesin-2 motor. SDS-PAGE analysis of the heterodimeric full-length FLA8/10 motor after FLAG-affinity-Tag purification of the C-terminally Flag-tagged FLA8 and the C-terminally 6X-His-tagged FLA10 subunits. **M**: marker. The C-terminally Flag-tagged FLA8 co-precipitates the 6X-His-tagged FLA10. The identity of the protein bands cannot be distinguished due to their similar sizes (∼87 kDa) (first lane after marker). The FLA10 subunit was N-terminally SNAP-tagged to distinguish the respective subunits. The C-terminally Flag-tagged FLA8 efficiently co-precipitated the 6X-His-tagged FLA10 subunit demonstrating the heterodimerization of the FLA8 and FLA10 subunits (second lane after marker). **(B) FLA8, FLA10 and KAP form a heterotrimeric kinesin-2 motor.** SDS-PAGE analysis of the full-length heterotrimeric FLA8/10/KAP (first lane after marker) and heterodimeric FLA8/10 (second lane after marker) kinesin-2 complexes after FLAG-affinity-Tag purification via the C-terminally Flag-tagged FLA8 subunit.

**Figure S2:**
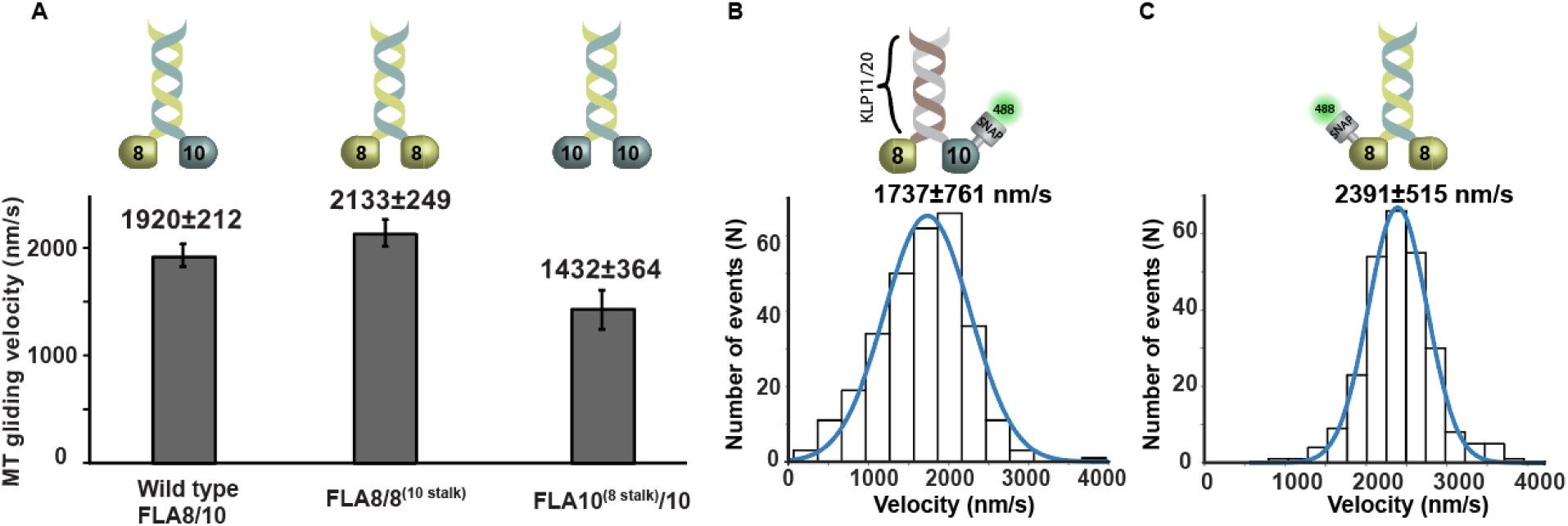
(A) The heterodimeric FLA8/10 kinesin-2 motor combines a fast FLA8 and a slow FLA10 head domain. Multiple-motor filament gliding assay of surface-attached motors moving fluorescently labeled microtubules (Suppl. movie 1). The wild type FLA8/10 (left) displays fast filament gliding velocities (1920±212 nm/s, N=30), as surface-binding via the stalk/tail abolishes the auto-inhibition (compare to the single-molecule behavior in Figure 2A). The chimeric FLA8/8 (middle) moves microtubules faster (2133±249 nm/s, N=30) than wild-type, whereas the FLA10/10 chimera (right) moves microtubules slower (1432±364 nm/s, N=30) than wild-type. Schematic (middle top) depicts the chimeric FLA8/8^(10 stalk)^ motor that was constructed by replacing the FLA10 head domain with the FLA8 head; similar strategy used for FLA10/10^(8 stalk)^ (right top). **(B) Catalytic heads as the main determinants of the kinesin-2-driven IFT velocity in *C. reinhardtii***. Schematic showing replacement of the head domains in the wild type KLP11/20 heterodimer with the FLA10 and FLA8 head domains creates a chimeric motor, FLA8^(KLP20)^/FLA10^(KLP11)^ with substantially increased velocity 1737±761 nm/s (N=296). **(C) Fast and fully active FLA8/8^(10 stalk)^ motor.** Single FLA8/8^(10 stalk)^ motors were tracked via the fluorescently labeled SNAP-tag on the N-terminus of the FLA8 subunit. Removal of the FLA10 head domain abolished the slow velocity distribution (Figure 2A, v1) and in the FLA8/8 chimera displayed a fast velocity of 2391±515 nm/s (N=262) similar to the fast v2 of the wild type FLA8/10 (Fig. 2A). Error bars indicate mean ± s.d. Data is fitted to a Gaussian distribution (±width of distribution, from two independent experiments).

**Figure S3:**
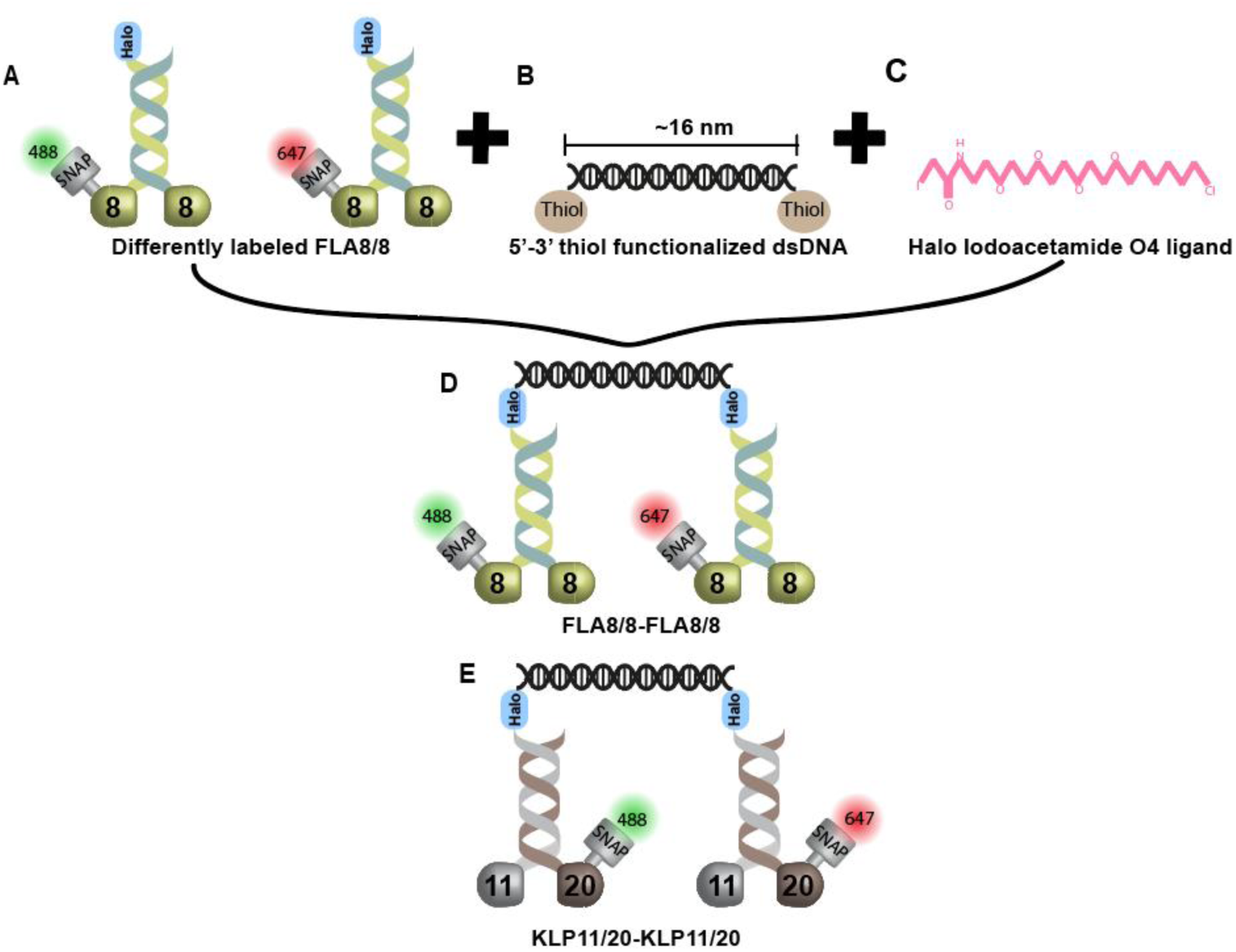
Coupling of two kinesin-2 motors with exclusive specificity. **(A)** The C-terminal Halo-tag was used to couple the Iodoacetamide-functionalized dsDNA to the differentially labeled FLA8/8 chimeras (SNAP^488^ and SNAP^647^ dyes). **(B)** dsDNA (48bp) was bi-functionalized with two thiol groups at both ends. **(C)** The Halo Iodoacetamide O4 ligand was coupled to the thiol-functionalized dsDNA. **(D)** Two FLA8/8 motors labeled with different fluorophores, ^SNAP488^FLA8^C-Halo^ /8^(10 stalk)^ and ^SNAP647^FLA8^C-Halo^/8^(10 stalk)^, were mixed in equimolar concentrations with the Iodoacetamide-bi-functionalized dsDNA to assemble a stable, covalently linked motor-DNA hybrid. **(E)** depicts coupling of two slow motors KLP11^C-Halo^/ ^SNAP488^20 and _KLP11_C-Halo_/_SNAP647_20._

**Figure S4:**
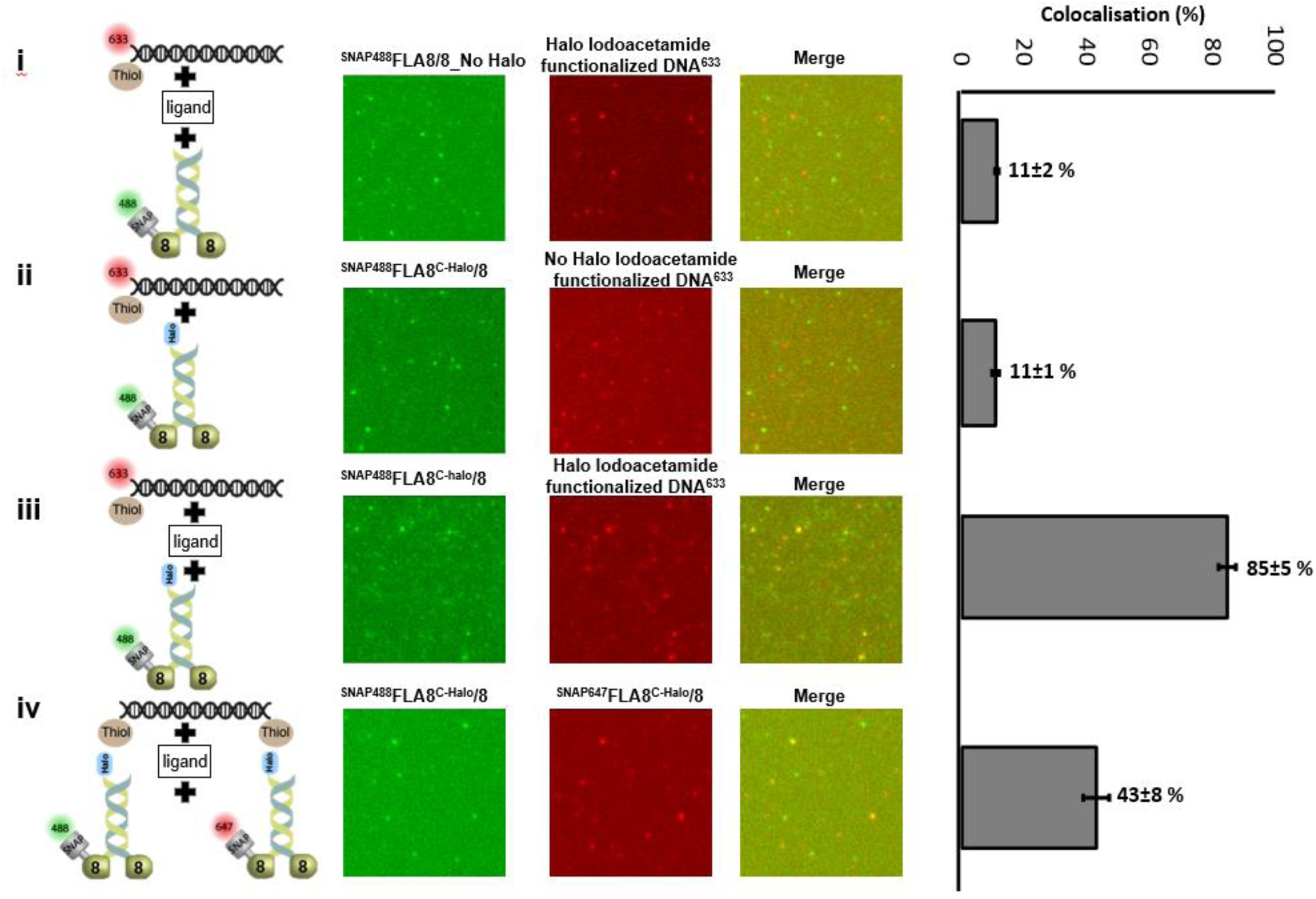
Highly specific DNA coupling and labeling of FLA8/8 motors. **i)** SNAP^488^-labeled FLA8/8 (green channel) motor that lacked the C-terminal Halo-tag was mixed in equimolar concentrations with the Atto-633-labeled and Iodoacetamide-functionalized dsDNA (red channel). Quantification on the right shows the corresponding colocalization efficiency of DNA-motor coupling *<u>in the absence of the C-terminal Halo-tag</u>* on the motor. **ii)** SNAP^488^-labeled and C-terminally Halo-tagged FLA8/8 motor was mixed in equimolar concentrations with the Atto-633-labeled and thiol-functionalized dsDNA that lacked the Iodoacetamide ligand (red channel). Quantification on the right shows the colocalization efficiency of DNA-motor coupling *<u>in the absence of the Iodoacetamide ligand</u>* on the dsDNA. **iii)** The presence of both the C-terminal Halo-tag on the FLA8/8 motor and the Iodoacetamide-functionalized dsDNA lead to robust colocalisation (85±5%). **iv)** SNAP^488^-labeled and C-terminally Halo-tagged FLA8/8 was (green channel) was mixed in equimolar concentrations with the SNAP^647^-labeled and C-terminally Halo-tagged FLA8/8 (red channel). The differently labeled motors were in turn mixed in equimolar concentrations with the Iodoacetamide-bi-functionalized (unlabeled) dsDNA. Quantification shows the 43±8% colocalisation efficiency, which is close to the statistically expected 50%. Numbers are mean±s.d.

**Figure S5:**
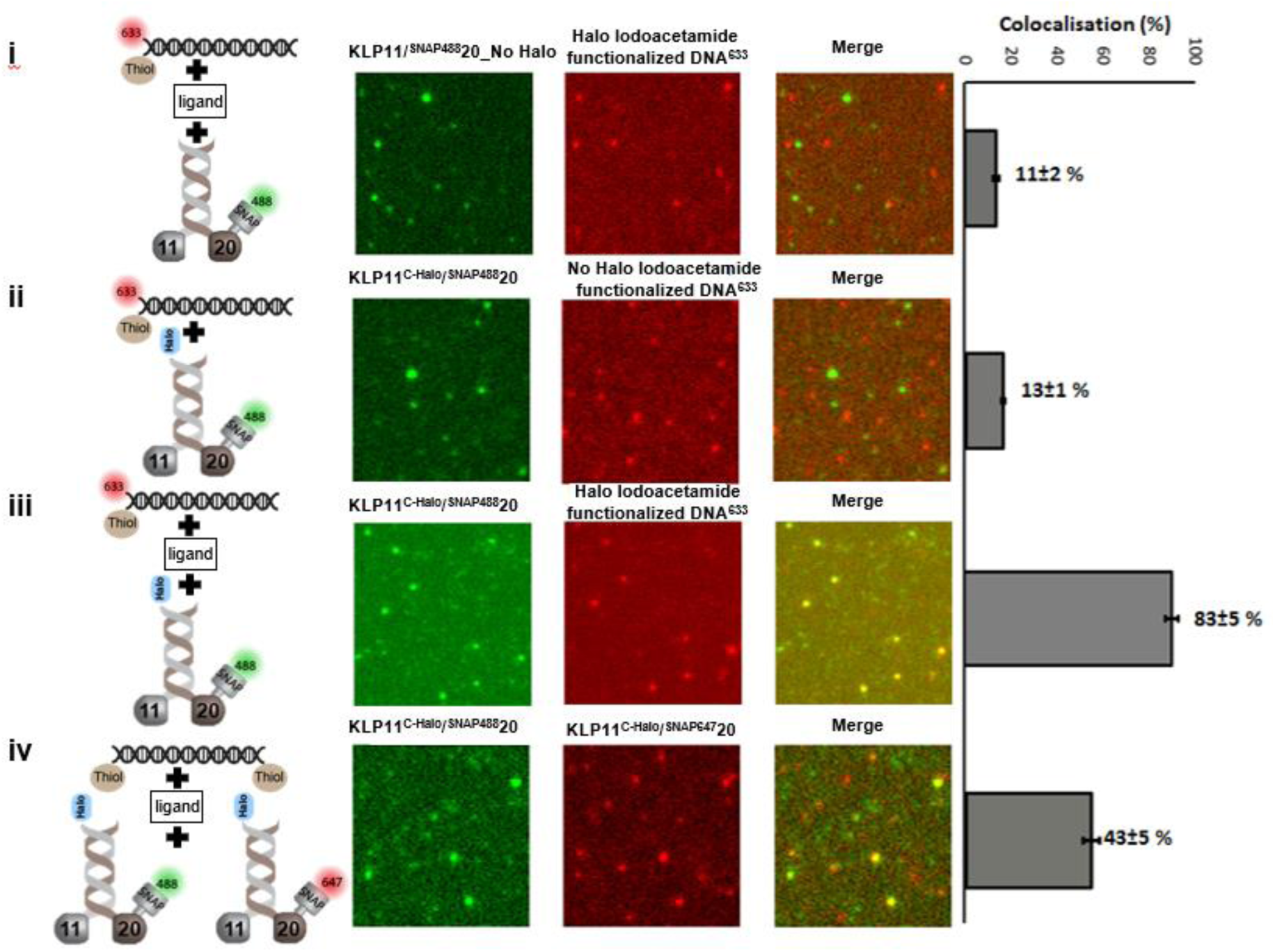
Highly specific DNA coupling and labeling of KLP11/20 motors. **i)** SNAP^488^-labeled KLP11/20 (green channel) motor that lacked the C-terminal Halo-tag was mixed in equimolar concentrations with the Atto-633-labeled and Iodoacetamide-functionalized dsDNA (red channel). Quantification on the right shows the corresponding colocalization efficiency of DNA-motor coupling *<u>in the absence of the C-terminal Halo-tag</u>* on the motor. **ii)** SNAP^488^-labeled and C-terminally Halo-tagged KLP11/20 motor was mixed in equimolar concentrations with the Atto-633-labeled and thiol-functionalized dsDNA that lacked the Iodoacetamide ligand (red channel). Quantification on the right shows the corresponding colocalization efficiency of the DNA-motor coupling *<u>in the absence of the Iodoacetamide ligand</u>* on the dsDNA. **iii)** The presence of both the C-terminal Halo-tag on the KLP11/20 motor and the Iodoacetamide-functionalized dsDNA lead to robust colocalisation (83±5%). **iv)** SNAP^488^-labeled and C-terminally Halo-tagged KLP11/20 (green channel) was mixed in equimolar concentrations with the SNAP^647^-labeled and C-terminally Halo-tagged KLP11/20 (red channel). The differently labeled motors were in turn mixed in equimolar concentrations with the Iodoacetamide-bi-functionalized (unlabeled) dsDNA. Quantification shows the 43±5% colocalisation efficiency, which is close to the statistically expected 50%. Numbers are mean±s.d.

**Figure S6:**
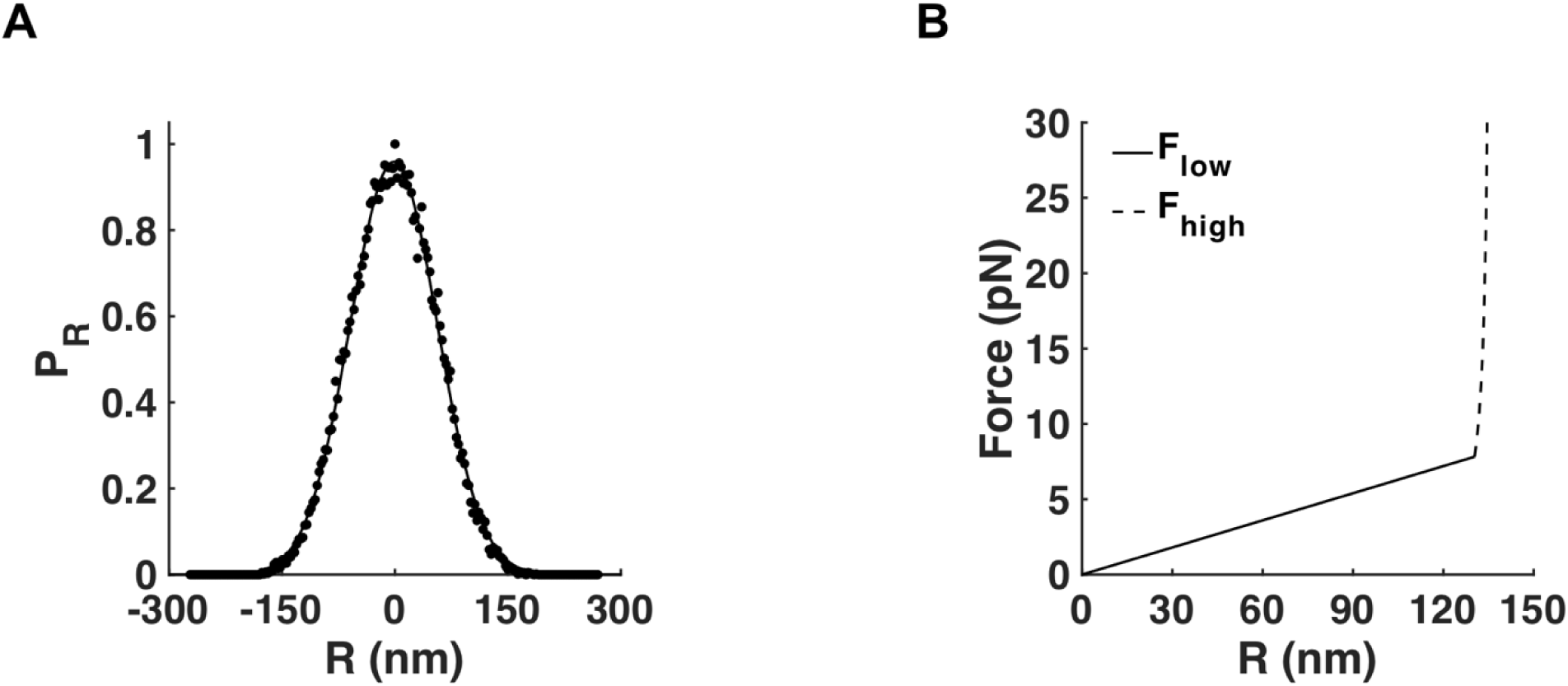
End-to-end distance distribution and force-extension curve used in the simulations. **(A)** Probability distribution of end-to-end distance of three-dimensional FJC model with a scaffold length of *136 nm*. The standard deviation of the distribution is *57.35 nm*. **(B)** The force-extension curve used in the simulations: at small forces, (*F* << *kT*/*l*; solid line), Hooke’s law is obeyed; at higher forces, (*F* >> *kT*/*l*; dashed line), forces increase exponentially with length (see Eq. (1)).

**Figure S7:**
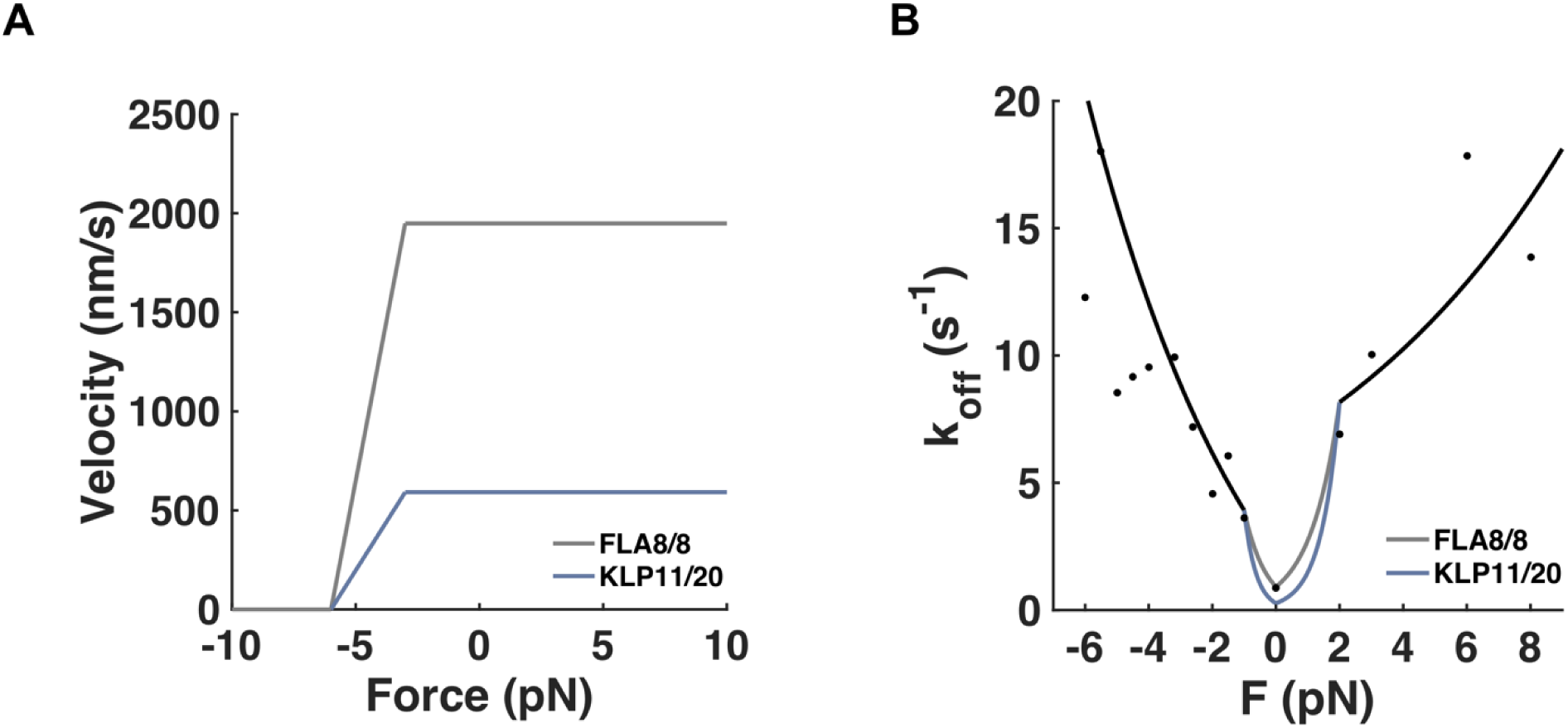
Model parameters. **(A)** Force-velocity curve of FLA8/8 (grey) and KLP11/20 (blue) used in the simulations. The force-velocity curve of KLP11/20 was estimated from [50]. The force-velocity curve of FLA8/8 was assumed to behave in the same manner as KLP11/20, but the unloaded velocity was adjusted from *592 nm/s* to *1949 nm/s*. **(B)** Force-dependent off-rates of FLA8/8 and KLP11/20 used in the simulations. Black circles show off-rate data measured experimentally by Milic et al. [46]. In our simulations, the off-rates of both FLA8/8 and KLP11/20 in the high-force regime were taken from exponential fits to the data points (black lines). The off-rates from exponential interpolations at low forces are shown for FLA8/8 (grey lines) and KLP11/20 (blue lines). The positive and negative forces refer to the assisting and hindering directions, respectively.

**Figure S8:**
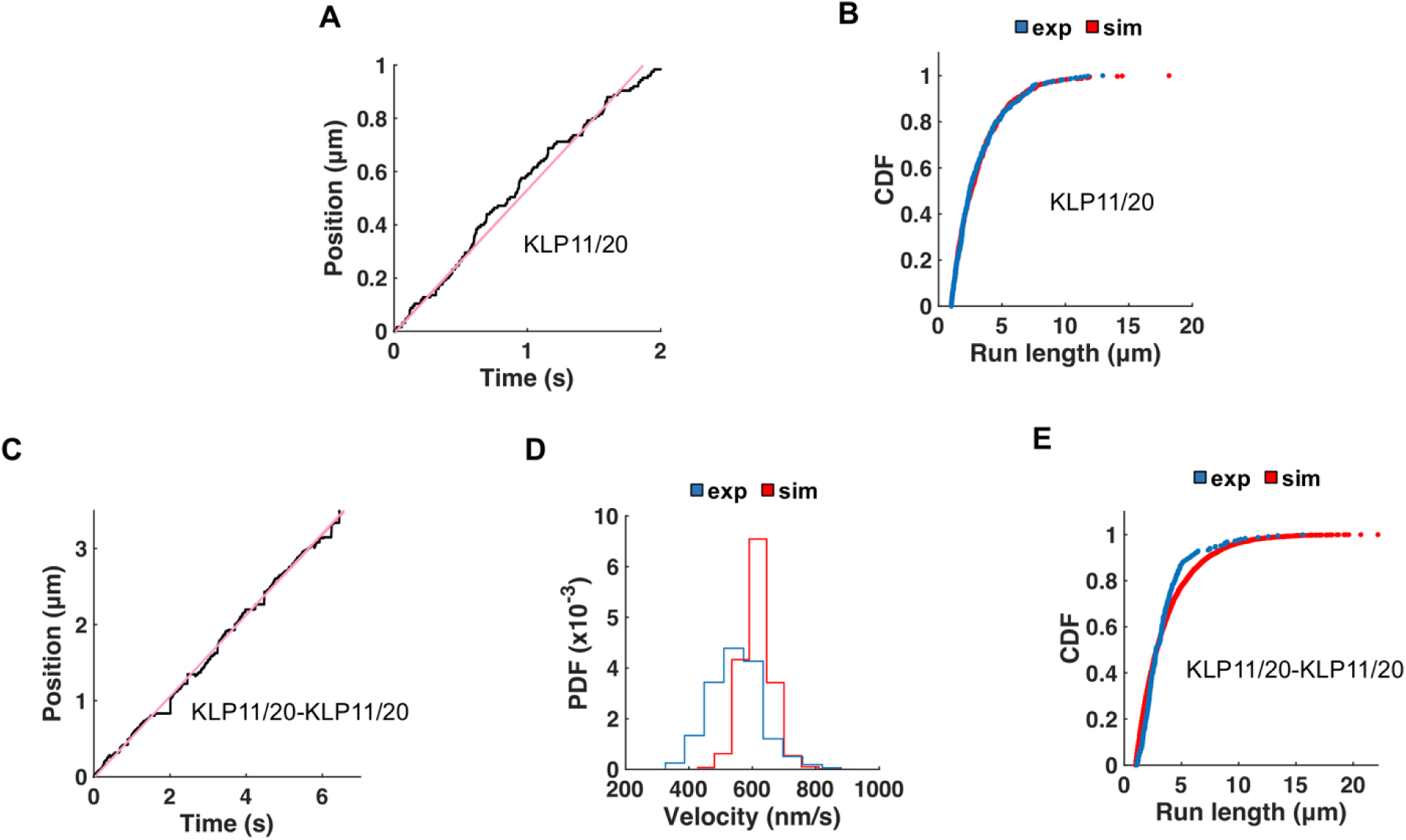
Velocities and run lengths of single KLP11/20 motor and KLP11/20-KLP11/20 co-transport. **(A)** Time-dependent positions (black) of single KLP11/20 motor from the simulations. Positions were traced at the motor heads for the single motor simulations. Mean velocity was calculated from the slope of each trace (red lines); values given in Table S1. **(B)** Cumulative distributions of experimental (*blue*) and simulated (*red*) run lengths of single KLP11/20 (Table S1). **(C)** Time-dependent positions (black) of KLP11/20-KLP11/20 co-transport from the simulations. The positions of the complexes were traced in the middle of the dsDNA. **(D)** Velocity distributions of KLP11/20-KLP11/20 from the experiment (blue) and simulation (red). The means and standard deviations were determined by fitting to a Gaussian function, resulting 513 ± 101 nm/s for the experiment and 586 ± 44 nm/s for the simulations, respectively. Simulated velocities were calculated from the slope of each trace (red lines in (C)). **(E)** The mean run lengths of KLP11/20-KLP11/20 co-transport were determined by fitting a CDF, resulting in 2.32 ± 0.03 μm for the experiment and 2.65 ± 0.09 μm for the simulation, respectively.

**Figure S9:**
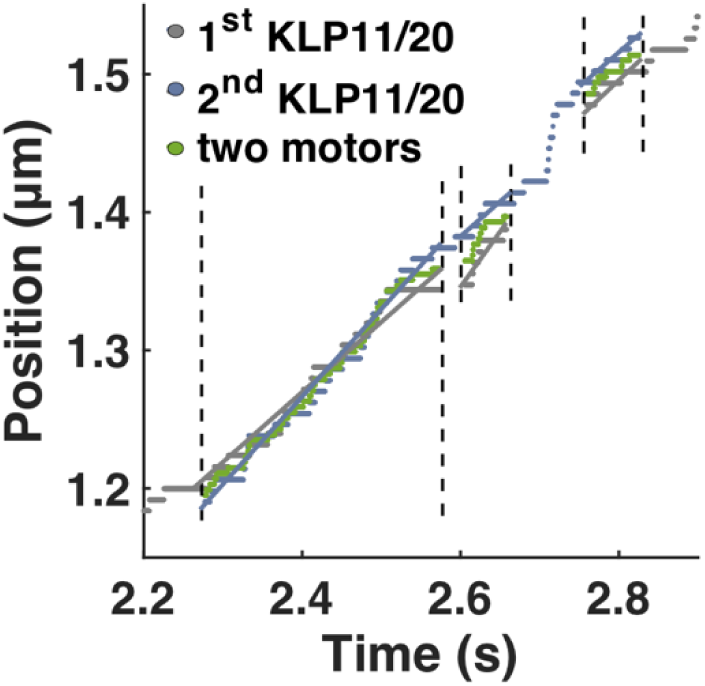
Simulated trace of KLP11/20-KLP11/20 co-transport. Traces show position of the first motor (grey) or the second motor (blue) when only one motor is bound to the microtubule, or the position of the middle of the complex (green) when both motors are simultaneously bound to the microtubule. The regions bounded by two vertical dashed lines represent periods of co-transport. The velocity shown in Figure 4F was calculated from the linear fits (as shown by the straight lines) to the trace during periods of co-transport.

## Supplementary Tables

**Table S1:** Summary of velocity and run length for experiments (Exp) and simulations (Sim). Values are mean ± s.d.

|  |  | FLA | FLA8/8-FLA8/8 | KLP | KLP11/20-KLP11/20 |
| --- | --- | --- | --- | --- | --- |
| Velocity (nm/s) | Exp | 1949 ± 408 | 1806 ± 339 | 592 ± 181 | 513 ± 101 |
|  | Sim | 1954 ± 117 | 1927 ± 124 | 587 ± 82 | 586 ± 44 |
| Run length (µm) | Exp | 2.12 ± 0.05 | 3.00 ± 0.06 | 2.18 ± 0.01 | 2.32 ± 0.03 |
|  | Sim | 2.12 ± 0.13 | 2.80 ± 0.11 | 2.28 ± 0.15 | 2.65 ± 0.09 |

**Table S2:** Summary of parameters used in the simulations. (*k*_off_(0) = velocity/run length).

| Variables | FLA8/8 | KLP11/20 |
| --- | --- | --- |
| $k_{on}$ (predicted) | 4 /s | 2.5 /s |
| $k_{off,0}$ | 0.9193 /s | 0.2716 /s |
| Unloaded velocity | 1949 nm/s | 592 nm/s |
| Stall force | 6 pN | 6 pN |
| $k_s$ | 0.06 pN/nm | 0.06 pN/nm |

## Supplementary movies

**Supplementary movie 1**: Multiple-motor gliding filament assay of wild type FLA8/10. The video is sped up 6X. Scale bar: 2 µm.

**Supplementary movie 2:** Colocalized assay of FLA8/8-FLA8/8 DNA hybrid. Top and middle channels represent individual motors tagged with SNAP^488^ and SNAP^647^ dyes respectively. The combined signal is shown in the bottom channel. The video is sped up 6X. Scale bar: 2 µm.

**Supplementary movie 3:** Colocalized assay of KLP11/20-KLP11/20 DNA hybrid. Top and middle channels represent individual motors tagged with SNAP^488^ and SNAP^647^ dyes respectively. The combined signal is shown in the bottom channel. The video is sped up 6X. Scale bar: 2 µm.

**Supplementary movie 4:** Movie shows a replication of an experimental movie of a FLA8/8-FLA8/8 DNA hybrid using data from the simulation. Model convolution is utilized with a Gaussian point spread function with a standard deviation of 300 nm. The top and middle channels represent individual motors tagged with SNAP^488^ and SNAP^647^ dyes, respectively. The combined signal is shown in the bottom channel. Scale bar: 2 µm.

**Supplementary movie 5:** Movie shows a replication of an experimental movie of a KLP11/20-KLP11/20 DNA hybrid using data from the simulation. Model convolution is utilized with a Gaussian point spread function with a standard deviation of 300 nm. The top and middle channels represent individual motors tagged with SNAP^488^ and SNAP^647^ dyes, respectively. The combined signal is shown in the bottom channel. Scale bar: 2 µm.

## References

1. Rosenbaum, J.L., and Witman, G.B. (2002). Intraflagellar transport. Nature reviews. Molecular cell biology 3, 813–825.

2. Scholey, J.M. (2003). Intraflagellar transport. Annual review of cell and developmental biology 19, 423–443.

3. Laib, J.A., Marin, J.A., Bloodgood, R.A., and Guilford, W.H. (2009). The reciprocal coordination and mechanics of molecular motors in living cells. Proceedings of the National Academy of Sciences of the United States of America 106, 3190–3195.

4. Cole, D.G., Diener, D.R., Himelblau, A.L., Beech, P.L., Fuster, J.C., and Rosenbaum, J.L. (1998). Chlamydomonas kinesin-II-dependent intraflagellar transport (IFT): IFT particles contain proteins required for ciliary assembly in Caenorhabditis elegans sensory neurons. The Journal of cell biology 141, 993–1008.

5. Badano, J.L., Mitsuma, N., Beales, P.L., and Katsanis, N. (2006). The ciliopathies: an emerging class of human genetic disorders. Annual review of genomics and human genetics 7, 125–148.

6. Nigg, E.A., and Raff, J.W. (2009). Centrioles, centrosomes, and cilia in health and disease. Cell 139, 663–678.

7. Pan, J., Wang, Q., and Snell, W.J. (2005). Cilium-generated signaling and cilia-related disorders. Laboratory investigation; a journal of technical methods and pathology 85, 452–463.

8. Scholey, J.M., and Anderson, K.V. (2006). Intraflagellar transport and cilium-based signaling. Cell 125, 439–442.

9. Anvarian, Z., Mykytyn, K., Mukhopadhyay, S., Pedersen, L.B., and Christensen, S.T. (2019). Cellular signalling by primary cilia in development, organ function and disease. Nature Reviews Nephrology 15, 199–219.

10. Bisgrove, B.W., and Yost, H.J. (2006). The roles of cilia in developmental disorders and disease. 133, 4131–4143.

11. Blacque, O.E., and Leroux, M.R. (2006). Bardet-Biedl syndrome: an emerging pathomechanism of intracellular transport. Cellular and molecular life sciences : CMLS 63, 2145–2161.

12. Pazour, G.J., and Rosenbaum, J.L. (2002). Intraflagellar transport and cilia-dependent diseases. Trends in cell biology 12, 551–555.

13. Kozminski, K.G., Johnson, K.A., Forscher, P., and Rosenbaum, J.L. (1993). A motility in the eukaryotic flagellum unrelated to flagellar beating. Proceedings of the National Academy of Sciences of the United States of America 90, 5519–5523.

14. Cole, D.G., Chinn, S.W., Wedaman, K.P., Hall, K., Vuong, T., and Scholey, J.M. (1993). Novel heterotrimeric kinesin-related protein purified from sea urchin eggs. Nature 366, 268–270.

15. Kozminski, K.G., Beech, P.L., and Rosenbaum, J.L. (1995). The Chlamydomonas kinesin-like protein FLA10 is involved in motility associated with the flagellar membrane. The Journal of cell biology 131, 1517–1527.

16. Pazour, G.J., Dickert, B.L., and Witman, G.B. (1999). The DHC1b (DHC2) isoform of cytoplasmic dynein is required for flagellar assembly. The Journal of cell biology 144, 473–481.

17. Porter, M.E., Bower, R., Knott, J.A., Byrd, P., and Dentler, W. (1999). Cytoplasmic dynein heavy chain 1b is required for flagellar assembly in Chlamydomonas. Molecular biology of the cell 10, 693–712.

18. Signor, D., Rose, L.S., and Scholey, J.M. (2000). Analysis of the roles of kinesin and dynein motors in microtubule-based transport in the Caenorhabditis elegans nervous system. Methods (San Diego, Calif.) 22, 317–325.

19. Signor, D., Wedaman, K.P., Rose, L.S., and Scholey, J.M. (1999). Two heteromeric kinesin complexes in chemosensory neurons and sensory cilia of Caenorhabditis elegans. Molecular biology of the cell 10, 345–360.

20. Wedaman, K.P., Meyer, D.W., Rashid, D.J., Cole, D.G., and Scholey, J.M. (1996). Sequence and submolecular localization of the 115-kD accessory subunit of the heterotrimeric kinesin-II (KRP85/95) complex. The Journal of cell biology 132, 371–380.

21. Signor, D., Wedaman, K.P., Orozco, J.T., Dwyer, N.D., Bargmann, C.I., Rose, L.S., and Scholey, J.M. (1999). Role of a class DHC1b dynein in retrograde transport of IFT motors and IFT raft particles along cilia, but not dendrites, in chemosensory neurons of living Caenorhabditis elegans. The Journal of cell biology 147, 519–530.

22. Mitchell, D.R. (2007). The evolution of eukaryotic cilia and flagella as motile and sensory organelles. Advances in experimental medicine and biology 607, 130–140.

23. Scholey, J.M. (2013). Kinesin-2: a family of heterotrimeric and homodimeric motors with diverse intracellular transport functions. Annual review of cell and developmental biology 29, 443–469.

24. Gilbert, S.P., Guzik-Lendrum, S., and Rayment, I. (2018). Kinesin-2 motors: Kinetics and biophysics. The Journal of biological chemistry 293, 4510–4518.

25. Mueller, J., Perrone, C.A., Bower, R., Cole, D.G., and Porter, M.E. (2005). The FLA3 KAP subunit is required for localization of kinesin-2 to the site of flagellar assembly and processive anterograde intraflagellar transport. Molecular biology of the cell 16, 1341–1354.

26. Sarpal, R., Todi, S.V., Sivan-Loukianova, E., Shirolikar, S., Subramanian, N., Raff, E.C., Erickson, J.W., Ray, K., and Eberl, D.F. (2003). Drosophila KAP interacts with the kinesin II motor subunit KLP64D to assemble chordotonal sensory cilia, but not sperm tails. Current biology : CB 13, 1687–1696.

27. Snow, J.J., Ou, G., Gunnarson, A.L., Walker, M.R., Zhou, H.M., Brust-Mascher, I., and Scholey, J.M. (2004). Two anterograde intraflagellar transport motors cooperate to build sensory cilia on C. elegans neurons. Nature cell biology 6, 1109–1113.

28. Teng, J., Rai, T., Tanaka, Y., Takei, Y., Nakata, T., Hirasawa, M., Kulkarni, A.B., and Hirokawa, N. (2005). The KIF3 motor transports N-cadherin and organizes the developing neuroepithelium. Nature cell biology 7, 474–482.

29. Brunnbauer, M., Mueller-Planitz, F., Kosem, S., Ho, T.H., Dombi, R., Gebhardt, J.C., Rief, M., and Okten, Z. (2010). Regulation of a heterodimeric kinesin-2 through an unprocessive motor domain that is turned processive by its partner. Proceedings of the National Academy of Sciences of the United States of America 107, 10460–10465.

30. Pan, X., Ou, G., Civelekoglu-Scholey, G., Blacque, O.E., Endres, N.F., Tao, L., Mogilner, A., Leroux, M.R., Vale, R.D., and Scholey, J.M. (2006). Mechanism of transport of IFT particles in C. elegans cilia by the concerted action of kinesin-II and OSM-3 motors. The Journal of cell biology 174, 1035–1045.

31. Stepp, W.L., Merck, G., Mueller-Planitz, F., and Okten, Z. (2017). Kinesin-2 motors adapt their stepping behavior for processive transport on axonemes and microtubules. EMBO reports 18, 1947–1956.

32. Andreasson, J.O., Shastry, S., Hancock, W.O., and Block, S.M. (2015). The Mechanochemical Cycle of Mammalian Kinesin-2 KIF3A/B under Load. Current biology : CB 25, 1166–1175.

33. Tuma, M.C., Zill, A., Le Bot, N., Vernos, I., and Gelfand, V. (1998). Heterotrimeric kinesin II is the microtubule motor protein responsible for pigment dispersion in Xenopus melanophores. The Journal of cell biology 143, 1547–1558.

34. Yamazaki, H., Nakata, T., Okada, Y., and Hirokawa, N. (1995). KIF3A/B: a heterodimeric kinesin superfamily protein that works as a microtubule plus end-directed motor for membrane organelle transport. The Journal of cell biology 130, 1387–1399.

35. Stepanek, L., and Pigino, G. (2016). Microtubule doublets are double-track railways for intraflagellar transport trains. Science (New York, N.Y.) 352, 721–724.

36. Iomini, C., Babaev-Khaimov, V., Sassaroli, M., and Piperno, G. (2001). Protein particles in Chlamydomonas flagella undergo a transport cycle consisting of four phases. The Journal of cell biology 153, 13–24.

37. Prevo, B., Mangeol, P., Oswald, F., Scholey, J.M., and Peterman, E.J. (2015). Functional differentiation of cooperating kinesin-2 motors orchestrates cargo import and transport in C. elegans cilia. Nature cell biology 17, 1536–1545.

38. Jordan, M.A., Diener, D.R., Stepanek, L., and Pigino, G. (2018). The cryo-EM structure of intraflagellar transport trains reveals how dynein is inactivated to ensure unidirectional anterograde movement in cilia. Nature cell biology 20, 1250–1255.

39. Mohamed, M.A.A., Stepp, W.L., and Okten, Z. (2018). Reconstitution reveals motor activation for intraflagellar transport. Nature 557, 387–391.

40. Imanishi, M., Endres, N.F., Gennerich, A., and Vale, R.D. (2006). Autoinhibition regulates the motility of the C. elegans intraflagellar transport motor OSM-3. The Journal of cell biology 174, 931–937.

41. Cole, D.G. (1999). Kinesin-II, the heteromeric kinesin. Cellular and molecular life sciences : CMLS 56, 217–226.

42. Stock, M.F., Guerrero, J., Cobb, B., Eggers, C.T., Huang, T.G., Li, X., and Hackney, D.D. (1999). Formation of the compact confomer of kinesin requires a COOH-terminal heavy chain domain and inhibits microtubule-stimulated ATPase activity. The Journal of biological chemistry 274, 14617–14623.

43. Coy, D.L., Hancock, W.O., Wagenbach, M., and Howard, J. (1999). Kinesin’s tail domain is an inhibitory regulator of the motor domain. Nature cell biology 1, 288–292.

44. Friedman, D.S., and Vale, R.D. (1999). Single-molecule analysis of kinesin motility reveals regulation by the cargo-binding tail domain. Nature cell biology 1, 293–297.

45. Ou, G., Blacque, O.E., Snow, J.J., Leroux, M.R., and Scholey, J.M. (2005). Functional coordination of intraflagellar transport motors. Nature 436, 583–587.

46. Milic, B., Andreasson, J.O.L., Hogan, D.W., and Block, S.M. (2017). Intraflagellar transport velocity is governed by the number of active KIF17 and KIF3AB motors and their motility properties under load. Proceedings of the National Academy of Sciences of the United States of America 114, E6830–e6838.

47. England, C.G., Luo, H., and Cai, W. (2015). HaloTag technology: a versatile platform for biomedical applications. Bioconjugate chemistry 26, 975–986.

48. Urh, M., and Rosenberg, M. (2012). HaloTag, a Platform Technology for Protein Analysis. Current chemical genomics 6, 72–78.

49. Arpag, G., Norris, S.R., Mousavi, S.I., Soppina, V., Verhey, K.J., Hancock, W.O., and Tuzel, E. (2019). Motor Dynamics Underlying Cargo Transport by Pairs of Kinesin-1 and Kinesin-3 Motors. Biophys J 116, 1115–1126.

50. Brunnbauer, M.D. (2012). Mechanische Untersuchungen heterodimerer Kinesin-2 Motoren. PHD thesis.

